# Specific inflammatory and mitochondrial signatures characterise the WSB/EiJ mice diet-induced obesity resistance capacity

**DOI:** 10.1101/504308

**Authors:** Jérémy Terrien, Isabelle Seugnet, Bolaji Seffou, Maria J. Herrero, James Bowers, Lamis Chamas, Stéphanie Decherf, Evelyne Duvernois-Berthet, Chakib Djediat, Bertrand Ducos, Barbara A. Demeneix, Marie-Stéphanie Clerget-Froidevaux

## Abstract

Energy balance disruption due to excess of food is considered to be one of the major players in the current worldwide obesity pandemic. In rodents, a high fat diet (HFD) induces not only obesity, but also inflammation and mitochondrial dysfunctions. To identify factors underlying diet-induced obesity (DIO) resistance we compared the wild-derived mouse strain WSB/EiJ, characterized by a striking resistance to DIO, with the more DIO-sensitive C57BL/6J strain. We analysed circulating levels of lipids, cytokines and adipokines as well as hypothalamic markers of inflammatory status and mitochondrial activity in both strains exposed to HFD for three days (3d) or eight weeks (8wk). To identify hypothalamic genes potentially involved in these differential regulations, we analysed the expression levels of 86 genes related to inflammation and mitochondrial pathways by high throughput microfluidic qPCR on RNA extracted from hypothalamic nuclei of the two strains of mice, under the different HFD treatments. After 3d and 8wk HFD, C57BL/6J mice, in contrast to WSB/EiJ, displayed significantly increased body weight gain, circulating levels of leptin, cholesterol, HDL and LDL. WSB/EiJ mice displayed a lower inflammatory status, both peripherally (lower levels of circulating cytokines) and centrally (less activated microglia in the hypothalamus) as well as more reactive mitochondria in the hypothalamus. Principal Component Analysis and gene ontology analysis of gene expression data allowed identifying the metabolic pathways involved. Strain-specific differential expression of several individual hypothalamic genes as well as differential effects of HFD between strains reinforced these results. Thus, adaptation to metabolic stress in the DIO-resistant WSB/EiJ strain implicates enhanced lipid metabolism, lower peripheral and hypothalamic inflammatory status and higher mitochondrial activity than in the C57BL/6J strain. These results point to the involvement of the hypothalamic inflammatory and mitochondrial pathways as key factors in the control of energy homeostasis and the resistance to DIO.

**Highlights:**

- Diet-induced-obesity resistance of WSB/EiJ implicates enhanced lipid metabolism
- WSB/EiJ mice are protected from HFD-induced peripheral and central inflammation
- Enhanced hypothalamic transport and signalling of endocrine molecules in WSB/EiJ
- WSB/EiJ hypothalamic mitochondria dynamics differ from C57BL/6J
- Differential regulation of hypothalamic inflammatory/mitochondria genes in WSB/EiJ

## 1. INTRODUCTION

The incidence of obesity and metabolism-related diseases such as type-2 diabetes, hypertension, and atherosclerosis, is inexorably increasing worldwide. Hence there is an urgent need to better understand the molecular mechanisms underlying physiological responses to changing calorific input.

The hypothalamus is the main central regulator of energy homeostasis [1]. In particular, two hypothalamic nuclei are involved in the maintenance of energy balance: the arcuate nucleus (ARC), and the paraventricular nucleus (PVN). The ARC integrates peripheral signals regarding metabolic status of the organism, such as insulin, leptin or other hormones, and thus monitors the energy status of the entire organism. The ARC stimulates melanocortinergic pathways in the PVN, which activate or repress food intake and energy expenditure [2]. Several studies have shown that high fat diet (HFD) in rodents initiates hypothalamic inflammation, especially in the ARC [3–5]. Just one to three days of HFD are sufficient to induce activation of both microglia and astrocytes leading to increased inflammatory markers in the ARC [6, 7]. This hypothalamic inflammation occurs prior to the onset of weight gain and peripheral inflammation, suggesting a neuroprotective response induced by hypothalamic nutrient sensing, rather than adipokines signalling to the hypothalamus. However, under a prolonged HFD, low-grade peripheral inflammation occurs, concomitant to weight gain, and hypothalamic inflammation and gliosis become established with continued HFD exposure [7]. Another important player in the central regulation of metabolism is the mitochondrion. Mitochondria have been long known to play major roles in cellular metabolism, but recently, mitochondrial dysfunction has also been linked to metabolic disorders [8]. In particular, in the hypothalamus, mitochondrial dynamics could participate in the control of food intake and energy expenditure [9]. Horvath and collaborators have shown that regulation of mitochondrial fusion by Mfn1 and Mfn2 is fundamental to the activation of NPY–AgRP neurons following exposure to a high-fat diet [10]. Moreover, mitochondrial fusion regulates neuronal firing via modulation of intracellular ATP levels in a mouse model of diet-induced obesity (DIO) [11].

Comparative phenotypic studies in mouse strains with differential susceptibility to DIO can provide perspectives into the pathophysiology of metabolic disorders. An interesting model is the wild-derived WSB/EiJ strain which is resistant to DIO, in contrast to C57BL/6J, an extensively studied laboratory strain prone to DIO and insulin resistance [12]. However, apart from the study by Lee and collaborators, highlighting important differences in insulin secretion and sensitivity between both strains, no other studies provide clues to understand the extraordinary DIO-resistance and healthy phenotype under HFD of WSB/EiJ. In this context, we hypothesized that the DIO-resistance observed in WSB/EiJ mice could be centrally regulated via the hypothalamus. Then, protective mechanisms favouring transport and signalling of endocrine signals could allow for rapid metabolic responses in hypothalamic areas, which should be reflected at the peripheral levels as compared to the C57BL/6J strain. To gain insight into which immediate mechanisms underlie the early onset response contributing to obesity resistance, we challenged DIO-resistant WSB/EiJ mice with a HFD treatment (for 3 days or 8 weeks) and compared their response with obesity-prone C57BL/6J mice. We focused our analysis on inflammatory markers (hypothalamic and peripheral), circulating lipids and adipokines, as well as hypothalamic expression of genes related to inflammation and mitochondrial pathways. After only three days of HFD, strain differences were observed in both central and peripheral responses involving inflammation, lipid fate and energy metabolism. This was accompanied by differential expression of hypothalamic genes involved in metabolism, inflammation and mitochondrial pathways.

## 2. MATERIALS AND METHODS

### 2.1. Animals and sampling

Animals were housed individually under a 12:12 light-dark cycle (07h00-19h00), maintained at 23°C, with food and drinking water provided *ad libitum*. Male C57BL/6J mice were provided by Charles River (l’Arbresle, France). WSB/EiJ breeders were purchased from The Jackson Laboratories (Maine, USA) and bred in house. At 4 weeks of age, i.e. at weaning, males were separated into 3 different diet groups to evaluate the responses to acute (3 days, 3d) and chronic (8 weeks, 8wk) exposure to a HFD *vs* a control diet (CTRL). Since then, CTRL and 8wk HFD animals were kept under their respective diets until 12 weeks of age. The 3d HFD group was also kept on the CTRL diet and then switched to HFD 3 days prior to 12 weeks of age. Control (D12450H) and HFD (D12451) diets contained 10% and 45% Kcal from fat respectively, and were matched for protein (20%) and sucrose (17%) contents (Research Diets Inc., Brogaarden, Gentofte, Denmark). Mice were weighed every week. At 12 weeks of age, retro-orbital blood was sampled just before mice were euthanized by decapitation. Trunk blood and tissue samples were collected in the morning. Blood samples were allowed to clot for 2 hours at room temperature (RT), centrifuged (20 min, 2 000 x g, RT) and serum supernatants stored at −20°C. Ependymal and inguinal white adipose tissues (eWAT, iWAT) were weighed before collection. All tissues were snap frozen in liquid nitrogen and stored at −80°C until further analysis. For immunohistochemistry and hypothalamic nuclei RNA extractions, whole brains were frozen for 1 minute in −35°C 2-methyl-butane (Isopentane, Sigma-Aldrich) and stored at −80°C. Animal experimentation protocols were validated by Ethics Committee n°68 regulated by the “Ministère de l’Enseignement Supérieur et de la Recherche” (France). Experimental procedures were authorised under reference numbers 68.032 and 00756.02.

### 2.2. Circulating metabolic parameters and serum cytokines

Serum adiponectin was assayed by ELISA (KMP0041, Life Technologies, US), according to manufacturer’s protocol. Leptin, resistin, C-Peptide 2, glucose-dependent insulinotropic peptide (GIP) and anorexigenic peptide PYY were simultaneously assayed on serum samples added with antiproteases (Complete EDTA Free, Roche, Meylan, France) using the Milliplex mouse metabolic hormone magnetic bead panel (MMHMAG-44K, Merck Millipore, Fontenay sous Bois, France). Circulating cytokine levels were analysed using the 10-spot V-plex kit Proinflammatory Panel 1 Mouse (Meso Scale Discovery, Rockville, USA). Data acquisition of multiplex analyses was performed at the Cochin Cytometry and Immunobiology Facility (France) using lectors Bioplex 200 (Biorad) and Sector Imager 2400 (Meso Scale Discovery). Lipid profile, hydroxybutyrate and total antioxidant status (TAS) were assayed on trunk serum using an Olympus AU-400 multiparametric analyzer at ‘Plateforme de Biochimie’ (INSERM U1149, France). Values for triglycerides, total cholesterol, high-density lipoprotein-cholesterol (HDL) and non-esterified fatty acids (NEFA) allowed the calculation of very low density lipoprotein-cholesterol (VLDL) and low density lipoprotein-cholesterol (LDL) following the Friedewald’s equation [13].

### 2.3. Hypothalamic immunohistochemistry

Frozen brains were post-fixed overnight (O/N) in paraformaldehyde (PFA) 4% in PBS, pH 7.4, immersed O/N in 30% sucrose in PBS and embedded in Frozen Section Compound (FSC 22R Clear LEICA Biosystems). Hypothalamic frozen coronal sections (30 μm) were processed using a cryostat (Leica CM 3050S cryostat, Leica Mannheim, Germany) and floating sections stored at −20°C in cryo-protectant solution until use. Sections were saturated in 10% normal goat serum (NGS) and 1% bovine serum albumin (BSA) in PBS for 1 hour at RT and incubated with rabbit primary antibody against microglia marker Iba-1 (1/200, Wako Chemicals, Japan) O/N at 4°C. Washed sections were incubated with the secondary anti-rabbit antibody Alexa Fluor 488 (1/500, Invitrogen) for 2 hours at RT. After washing, sections were incubated with the chicken primary antibody against GFAP (1/300, Abcam) O/N at 4°C, followed by the secondary anti-chicken antibody Alexa Fluor 594 (1/500 Invitrogen) for 2 hours at RT. DAPI staining was used as a nucleus marker. Sections were mounted onto SuperFrost/Plus glass slides with ProLong antifade (Invitrogen) and stored in darkness at 4°C. Slides were observed under SP5 Leica confocal microscopy at ImagoSeine Facility (Paris, France). Two independent observers quantified microglial density and microglial surface area (μm^2^) in the PVN and ARC regions of interest (ROI) using the Image J software.

### 2.4. Hypothalamic nuclei laser-capture microdissection and RNA extraction

Frozen brains were sectioned at a thickness of 30 μm using a cryostat. Cryosections were mounted onto PEN (Polyethylene naphtalate)-membrane glass slides (Leica Microsystems) and stored at −80°C with desiccant. They were lightly fixed in 75% ethanol just before rapid Cresyl violet staining (Sigma-Aldrich), and dehydration in a graded ethanol series followed by an under vacuum air-dry at RT for 5 minutes. All solutions were prepared with RNase-free water.

The microdissection was performed using a LEICA 6500 laser-capture microdissection system (Leica Mannheim, Germany) according to the manufacturer’s protocol, to isolate the PVN and the ARC of the hypothalamus. Samples were collected directly into RLT buffer (RNeasy Plus Micro Kit, Qiagen) added with DTT at 40 μM and immediately stored at −80°C. Total RNA samples were extracted using the RNeasy Plus Micro Kit (Qiagen) according to the manufacturer’s protocol to remove genomic contamination and stored at −80°C. All RNA samples were quantified on the Qubit 2.0 Fluorometer (Invitrogen life technologies) and RNA integrity was evaluated on the Agilent 2100 Bioanalyzer.

### 2.5. Reverse transcription and microfluidic qPCR

#### 2.5.1. qRT-PCR

Complementary DNA (cDNA) synthesis was performed using Reverse Transcription Master Mix from Fluidigm® according to the manufacturer’s protocol with random primers in a final volume of 5 μL containing 6 ng total RNA. cDNA samples were diluted by adding 20 μL of low TE buffer [10 mM Tris; 0.1 mM EDTA; pH = 8.0 (TEKNOVA)] and stored at −20°C. For specific target pre-amplification 1.25 μL of each diluted cDNA was used for multiplex pre-amplification with Fluidigm® PreAmp Master Mix at 19 cycles. In a total volume of 5 μL the reaction contained 1 μL of pre-amplification mastermix, 1.25 μL of cDNA, 1.25 μL of pooled TaqMan® Gene Expression assays (86 target genes related to inflammation and mitochondria pathways, and 10 housekeeping genes - see list in supplementary material - Life Technologies, ThermoFisher) with a final concentration of each assay of 180 nM (0.2X) and 1 μL of PCR water. The cDNA samples were subjected to pre-amplification following the temperature protocol: 95°C for 2 min, followed by 19 cycles at 95°C for 15 s and 60°C for 4 min. The pre-amplified cDNA were diluted 5X by adding 20 μL of low TE buffer (TEKNOVA) and stored at −20°C before qPCR. High-throughput real time PCR was performed on the qPCR-HD-Genomic Paris Centre platform, using the high-throughput platform BioMark™ HD System and the 96.96 GE Dynamic Arrays (Fluidigm). Six μL of sample master mix (SMM) consisted of 1.8 μL of 5X diluted pre-amplified cDNA, 0.3 μL of 20X GE Sample Loading Reagent (Fluidigm) and 3 μL of TaqMan® Gene Expression PCR Master Mix (ThermoFisher). Each 6 μL assay master mix (AMM) consisted of 3 μL of TaqMan® Gene Expression assay 20X (ThermoFisher) and 3 μL of 2X Assay Loading Reagent (Fluidigm). Five μL of each SMM and each AMM premixes were added to the dedicated wells. The samples and assays were mixed inside the chip using HX IFC controller (Fluidigm). Thermal conditions for qPCR were: 25°C for 30 min and 70°C for 60 min for thermal mix; 50°C for 2 min and 95°C for 10 min for hot start; 40 cycles at 95°C for 15 s and 60°C for 1 min.

#### 2.5.2. Microfluidic qPCR data analysis

data analysis was done using Fluidigm Real-time PCR Software (version 4.1.3), with manual determination of the fluorescence threshold (CT), the quality threshold which was set by default (0.65) and Linear (Derivative) as a baseline correction. The three most stable housekeeping genes in PVN and ARC were selected from 10 housekeeping genes chosen from the bibliography using SlqPCR package (version 1.42.0) based on Vandesompele et al method [14]. A custom R tool was constructed to measure relative gene expression levels according to the ddCT method (DCT, DDCT, Fold Change (FC)) [15] and to perform statistical tests (one- and two-way non-parametric ANOVA with permutation). This was done by using the following R packages: RVAideMemoire (version 0.9-61), ez (version 4.4-0) and coin (version 1.1-3). DCT values were used for one- and two-way ANOVA with FDR correction (p<0.05) and respectively, 100000 permutations / 1000 permutations post-tests and 1000 permutations / 10000 permutations post-tests. Graphical representations were performed using FC values on GraphPad Prism (version 6.01). Outliers were removed using Dixon test from the Outliers package in R (version 0.14).

### 2.6. Bioinformatic data analysis

A custom R tool was designed to perform Heatmap using R packages: gplots (version 3.0.1), R-color-Brewer (version 1.1-2), and HEATMAP.2 function. Fold Change values for mitochondria and inflammation genes were used and transformed to log2 [16]. Scale was applied manually for each gene (horizontally) using SCALE function in R and a row dendrogram was performed. Color breaks were defined manually to have a color transition from green (low expression), yellow (mean expression) to red (high expression). Gene functional analysis (pathway) was identified using the bioinformatics enrichment tool DAVID (version 6.8).

### 2.7. Statistical analyses

Unless specifically mentioned, all data are presented as medians with ranges (GraphPad Prism 6 software). Unless for qPCR data analysis (see above), statistical-parametric tests were applied by using linear model (after data transformation when necessary), followed by post-hoc analysis (in case of normal distribution and homoscedasticity) with the R software 3.1.3. When data normalisation was not possible or when group numbers were small (for electron microscopy and immunohistochemistry analyses), non-parametric permutation tests were performed using the ‘lmp’ or ‘aovp’ functions (package ‘LmPerm’ 2.1.0). These analyses were run on all samples as a function of diet, strain, diet:strain and time (when appropriate), Data correction for body weight was applied when convenient according to Speakman et al [17]. Data sets are either a pool or representative of 3 independent experiments with n=6 animals / group (except for confocal image analysis n=3-4). To pool replicates of cytokine results, normalisation between assays was required. Dixon’s Q test was used for identification and rejection of outliers. Significant differences in graphs are indicated by different letters on top of bars. The level of significance was set at P-values ≤ 0.05.

## 3. RESULTS

### 3.1. Resistance to DIO in WSB/EiJ mice is associated with increased lipid metabolism

To gain insights into which mechanisms underlie the obesity-resistant response in the early stages of obesogenic conditions, we challenged WSB/EiJ mice to a HFD treatment and compared their response to obesity-prone C57BL/6J mice. WSB/EiJ mice displayed a strong resistance to the obesogenic treatment, as the HFD challenge did not alter body weight variations in comparison to control animals, regardless of challenge duration (3d or 8wk) (Fig. 1A-B and Table 1). In contrast, in C57BL/6J mice, HFD induced an increase in body weight (BW) gain within 3 days, which was confirmed after 8wk challenge (Fig1A-B and Table 1). At the end of treatment, the 8wk group displayed a 6.7% BW increase compared to control animals. This resulted in strong differences between WSB/EiJ mice and their C57BL/6J counterparts at the end of the 12-week experiment (Fig.1A and Table 1). Accordingly, the proportion of ependymal white adipose tissue (% eWAT) was not altered by HFD in WSB/EiJ mice (Fig.1C), while C57BL/6J displayed a 120% increase after 8 weeks of challenge. No significant variation was found in inguinal WAT (% iWAT) in either strain (Fig.1D).

**Table 1.** Summary table of the statistics (F and p-values) for parameters representative of body composition (related to Figure 1), circulating lipids (related to Figure 2), metabolic hormones (related to Figure 3) and inflammatory hormones (related to Figure 4). Statistical parametric tests were applied by using linear model (after data transformation when necessary), followed by post-hoc analysis. Analyses were run on all samples as a function of Strain, Diet and Strain*Diet, correcting data to body weight when convenient. Significant differences are indicated by different letters to account for between strains and between diet groups differences.

|  | Parameter | Strain |  | Diet |  | Strain*Diet |  | Posthoc |
| --- | --- | --- | --- | --- | --- | --- | --- | --- |
|  |  | F-value | p-value | F-value | p-value | F-value | p-value | BI6 CTRL-3D-8WK; WSB CTRL-3D-8WK |
| **Body composition**<br>(Figure 1) | *Final BW* | 1894 | < 0.0001*** | 8.0 | 0.0005*** | 5.5 | 0.005** | a - ab - b ; c - c - c |
|  | *Incremental BW* | 5000 | < 0.0001*** | 5000 | < 0.0001*** | 5000 | < 0.0001*** | a - b - c ; d - ad - d |
|  | *% eWAT* | 142 | < 0.0001*** | 36 | < 0.0001*** | 17.9 | < 0.0001*** | a - b - c ; d - ad - ad |
|  | *% iWAT* | 5.0 | 0.03* | 5.2 | 0.008** | 0.4 | 0.65 | a - ab - ab ; ab - b - b |
| **Circulating lipids**<br>(Figure 2) | *HDL* | 14.3 | 0.0004*** | 48.6 | < 0.0001*** | 5.8 | 0.005** | ac - bd - b ; a - c - cd |
|  | *Cholesterol* | 26.3 | < 0.0001*** | 22.3 | < 0.0001*** | 6.0 | 0.004** | a - b - b ; a - a - a |
|  | *LDL* | 66.8 | < 0.0001*** | 18.9 | < 0.0001*** | 13.1 | < 0.0001*** | a - b - b ; a - a - a |
|  | *Triglycerides* | 4.1 | 0.05* | 0.8 | 0.47 | 0.7 | 0.51 | / |
|  | *VLDL* | 4.1 | 0.05* | 0.8 | 0.47 | 0.7 | 0.51 | / |
|  | *NEFA* | 16.5 | 0.0002*** | 0.62 | 0.54 | 0.29 | 0.75 | ab - ab - a ; b - ab - ab |
|  | *Hydroxybotyrate* | 189 | < 0.0001*** | 5.3 | 0.007** | 6.5 | 0.003** | a - a - a ; b - c - bc |
| **Metabolic hormones**<br>(Figure 3) | *Leptin* | 47.2 | < 0.0001*** | 10.2 | 0.0002*** | 12.1 | < 0.0001*** | a - bd - b ; a - c - ad |
|  | *Adiponectin* | 4.6 | 0.04* | 10.1 | 0.0002*** | 0.2 | 0.83 | ab - a - ab ; b - a - b |
|  | *Resistin* | 1.7 | 0.20 | 1.0 | 0.37 | 7.8 | 0.001** | a - b - ab ; ab - a - ab |
|  | *C-peptide 2* | 1.5 | 0.23 | 0.7 | 0.49 | 9.7 | 0.0003*** | a - b - ab ; b - ab - ab |
| **Inflammatory hormones**<br>(Figure 4) | *IFN $\gamma$* | 243 | < 0.0001*** | 1.5 | 0.23 | 2.3 | 0.11 | a - a - a ; b - b - b |
|  | *KC/GRO* | 68.8 | < 0.0001*** | 0.05 | 0.95 | 0.40 | 0.68 | a - a - a ; b - b - b |
|  | *IL-5* | 93.1 | < 0.0001*** | 4.7 | 0.01* | 1.2 | 0.31 | a - a - a ; b - b - b |
|  | *IL-10* | 198 | < 0.0001*** | 0.52 | 0.60 | 1.0 | 0.37 | a - a - a ; b - b - b |
|  | *IL-6* | 44.3 | < 0.0001*** | 0.17 | 0.84 | 0.66 | 0.52 | a - a - a ; b - b - b |
|  | *TNF- $\alpha$* | 1.0 | 0.32 | 0.08 | 0.92 | 0.54 | 0.59 | / |
|  | *IL-1 $\beta$* | 43.0 | < 0.0001*** | 1.2 | 0.32 | 0.18 | 0.84 | ac - a - a ; b - bc - b |
|  | *TAS* | 78.7 | < 0.0001*** | 1.1 | 0.35 | 0.94 | 0.39 | a - a - a ; b - b - b |

**Figure 1:**
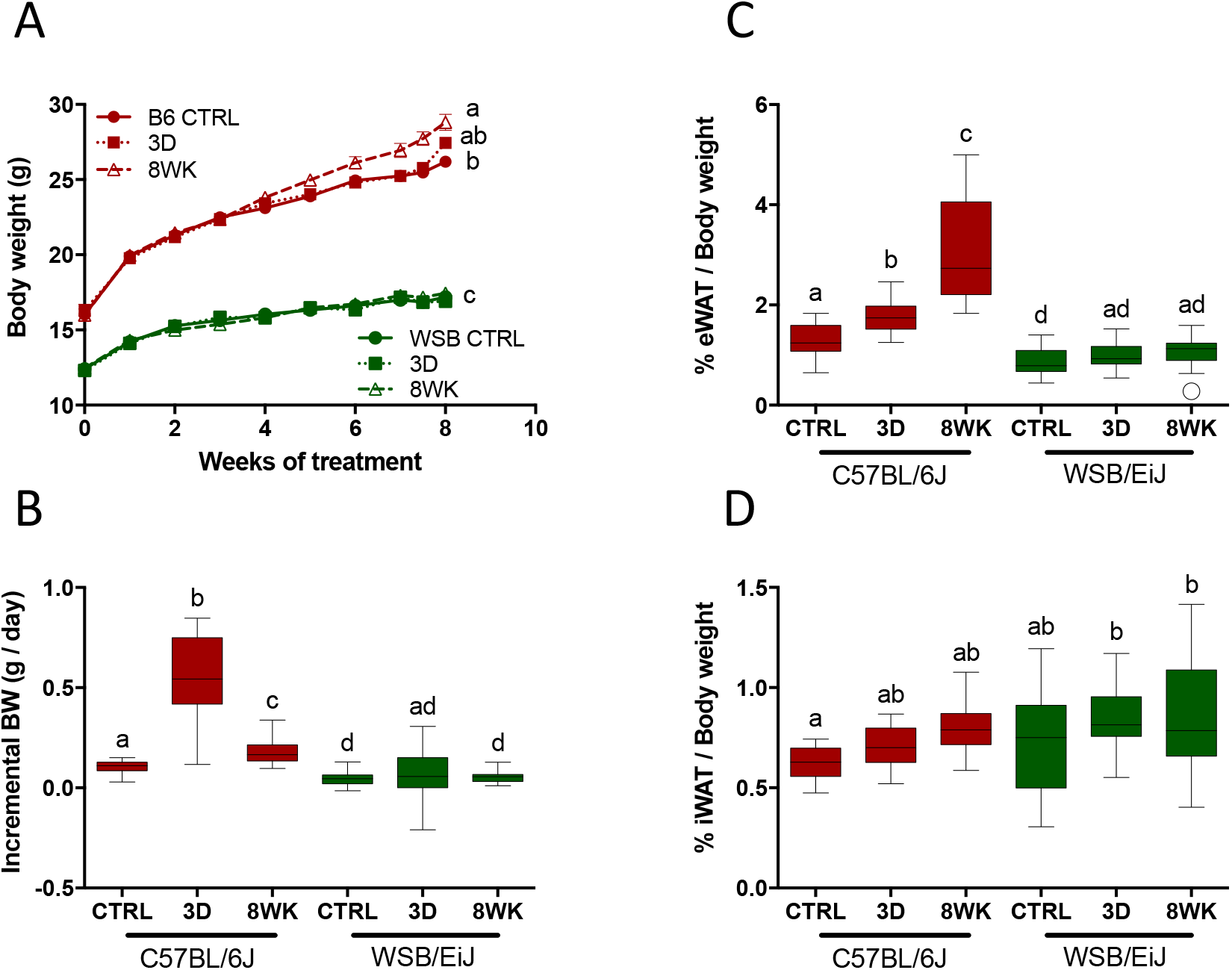
WSB/EiJ mice are resistant to HFD-induced obesity. Twelve week-old male C57BL/6J (B6) and WSB/EiJ (WSB) mice were experimented at weaning (4 weeks of age) during 8 weeks and were either maintained under control feeding (CTRL), or challenged with HFD for the last 3 days of the period (3D) or during the whole 8-week experiment duration (8WK). A. Growth curve upon weekly measurements (n=22-34 per group); B. Incremental body weight (BW in g/day) was calculated to account for the slope of the BW gain during the treatment ([Initial BW – Final BW] / nb of days) – the initial body weight was taken after 4 weeks of treatments for groups CTRL and 8WK, i.e. when differences between groups were observed; in group 3D, last 3 days of challenge only (before sampling) were considered (N=22-34 per group); C. Percentage of ependymal white adipose tissue (eWAT) measured at the end of the experiment (referred to final BW; N=14-18 per group); D. Percentage of inguinal white adipose tissue (iWAT) measured at the end of the experiment (referred to final BW; N=11-15 per group). Graphs represent medians using box and whiskers, except for growth curve (Mean ± SEM). Significant differences were indicated by different letters to account for between strain and between diet group differences (P≤0.05; GLM).

Next, we verified whether resistance to the obesogenic challenge in WSB/EiJ mice was reflected in circulating lipid parameters (Fig.2 and Table 1). While high-density lipoproteins (HDL), which are involved in cholesterol transport and removal, rapidly increased in both strains under HFD (Fig.2A), efficient export of cholesterol occurred only in WSB/EiJ mice. This effect was demonstrated by unchanged cholesterol levels in WSB/EiJ, whereas they increased in C57BL/6J mice (Fig.2B). Circulating cholesterol increased significantly by more than 30% after 3d treatment in C57BL/6J, and remained high after 8wk HFD (Fig.2B). Similarly, strain specific changes were seen for low density lipoproteins (LDL) with HFD-induced increases in C57BL/6J, but not in WSB/EiJ (Fig.2C). Triglyceride levels remained unchanged by HFD in both strains, indicating normal blood transfer of these main constituents of body fat (Fig.2D), consistent with unaltered levels of VLDL which are involved in transport of triglycerides (Table 1).

**Figure 2:**
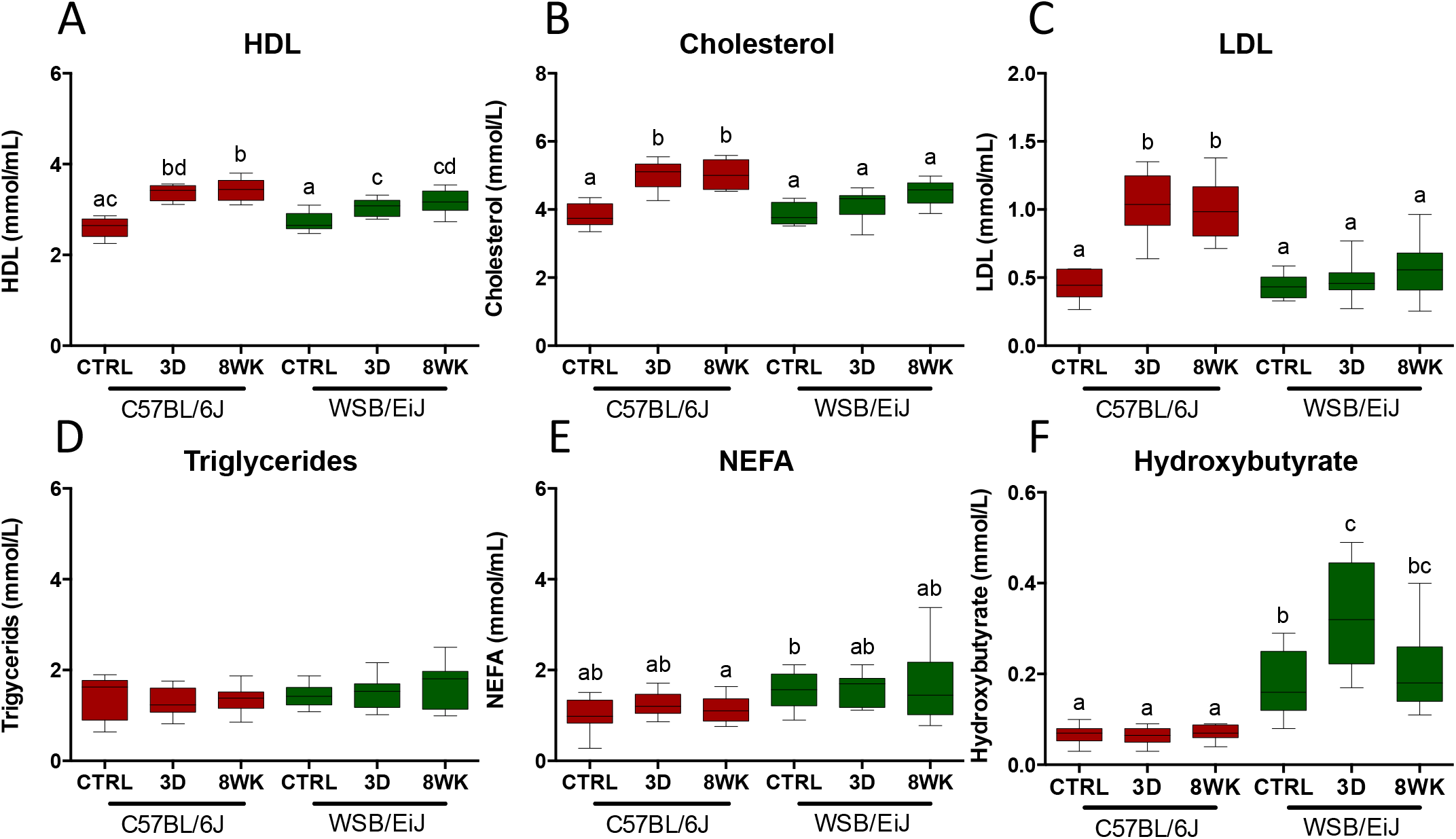
Patterns of circulating lipids are less altered under HFD in WSB/EiJ than in C57BL/6J mice. Compared analysis of circulating lipids in twelve week-old C57BL/6J and WSB/EiJ male mice either maintained under control feeding (CTRL), or challenged with HFD for the last 3 days of the period (3D) or during the whole 8-week experiment duration (8WK). A. High density lipoprotein (HDL in mmol/mL; N=10-12 per group); B. Total cholesterol (in mmol/L; N=10-12 per group); C. Low density lipoprotein (LDL in mmol/mL; N=8-11 per group); D. Triglycerides (in mmol/L; N=11-12 per group); E. Non-esterified fatty acids (NEFA in mmol/mL; N=10-12 per group); F. Hydroxybutyrate (in mmol/L; N=10-12 per group). Graphs represent medians using box and whiskers. Significant differences were indicated by different letters to account for between strain and between diet group differences (P≤0.05; GLM).

Non-esterified fatty acids (NEFA) levels also remained constant, unchanged by HFD in both strains (Table 1–Fig.2E). However, a higher variability was seen in the WSB/EiJ after 8wk HFD, suggesting enhanced hydrolysis of fats in this strain. Further supporting such hypothesis of a lipid-metabolising phenotype of WSB/EiJ strain, basal levels of hydroxybutyrate were higher in WSB/EiJ and sharply increased after 3d HFD, followed by a return to basal levels at 8wk, indicating enhanced β-oxidation as a fast response to lipid overload. In contrast, hydroxybutyrate levels were extremely low in C57BL/6J, and unaffected by the HFD challenge (Fig.2F and Table 1).

Levels of metabolic markers associated with appetite and lipid-glucose homeostasis were measured (Fig.3 and Table 1). While leptin, adiponectin and resistin are all produced by expanded adipocytes, C57BL/6J mice showed concomitant variations of leptin levels with the cumulative pattern of % eWAT under HFD. Three days of HFD triggered a marked increase of leptin in C57BL/6J mice, which was sustained through the 8wk HFD. In contrast, leptin levels remained low throughout the HFD challenge in WSB/EiJ mice (similar to the C57BL/6J control group), even transiently decreasing after 3d HFD (Fig.3A). Adiponectin was significantly regulated only in WSB/EiJ, reaching a peak at 3d but returning to basal levels at 8wk (Fig.3B). C57BL/6J showed the same trends, but variations were not significant. Resistin levels were differentially affected as a function of strain (Fig.3C and Table 1). In C57BL/6J mice, 3d HFD induced increased levels of resistin compared to controls, whereas no significant change was observed in WSB/EiJ mice. In addition, C-peptide 2, which is produced in equimolar concentrations with insulin, showed the same response trend as resistin (Fig.3D). HFD treatment provoked an increase of C-Peptide 2 levels at 3d in C57BL/6J mice, but returned to baseline by the end of the 8wk challenge. In contrast, WSB/EiJ mice displayed slightly higher basal levels of C-peptide 2 under control conditions. Similar strain-dependent inverted regulations were found in glucose-dependent insulinotropic peptide (GIP), but statistical analysis was inconclusive. Anorexigenic peptide PYY, released in response to feeding as a peripheral signal to reduce appetite, was at the limit of detection levels in both strains. However, in C57BL/6J a higher number of samples were above detection level when compared to WSB/EiJ (8 vs. 12 out of 18 for C57BL/6J and WSB/EiJ mice, respectively), suggesting enhanced appetite in WSB/EiJ strain (data not shown).

**Figure 3:**
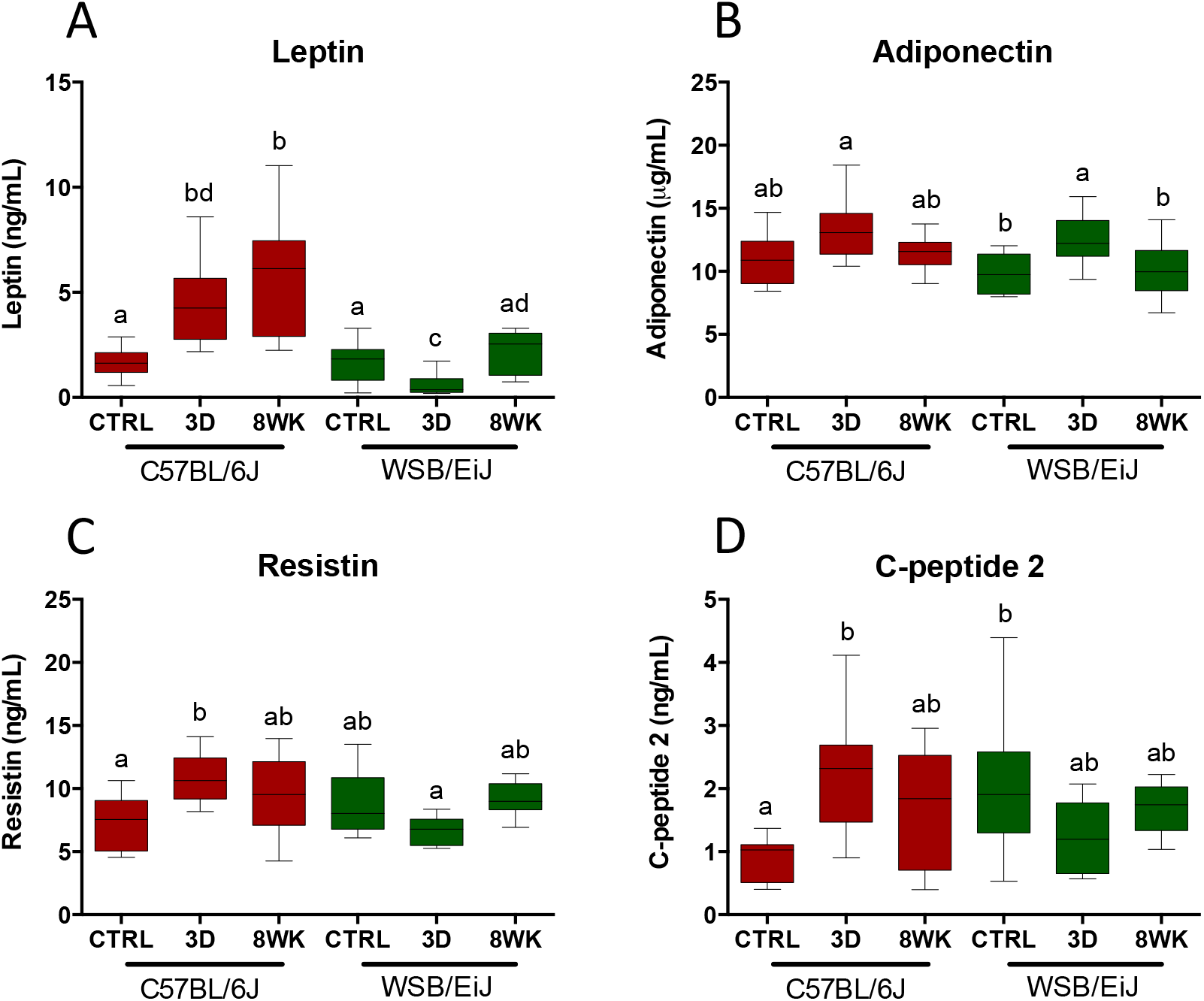
WSB/EiJ mice show differential regulations of circulating leptin levels and metabolic markers under HFD as compared to C57BL/6J strain. Compared analysis of main circulating biomarkers for energetic metabolism and glucose homeostasis in twelve week-old C57BL/6J and WSB/EiJ male mice either maintained under control feeding (CTRL), or challenged with HFD for the last 3 days of the period (3D) or during the whole 8-week experiment duration (8WK). A. Circulating leptin (in ng/mL; N=9-12 per group); B. Adiponectin (in μg/mL; N=9-11 per group); C. Resistin (in ng/mL; N=8-12 per group); D. C-peptide 2 (in ng/mL; N=8-12 per group). Graphs represent medians using box and whiskers. Significant differences were indicated by different letters to account for between strain and between diet group differences (P≤0.05; GLM).

### 3.2. WSB/EiJ mice are remarkably protected from peripheral and central inflammation under HFD

Circulating cytokines in response to HFD were compared between C57BL/6J and WSB/EiJ mice to investigate early occurrence of low-grade chronic inflammation associated with obesity (Fig.4 and Table 1). Overall, C57BL/6J displayed markedly higher baseline levels for most of the reported circulating cytokines including IFN-γ, KC/GRO, IL-5, IL-10 and IL-6 (Fig.4A-E), with the exception of TNF-α and IL-1β (Fig.4F-G). TNF-α levels displayed no strain effects (Fig.4F), whereas baseline levels of the anorexigenic and pyrogenic IL-1β were markedly higher in WSB/EiJ than in C57BL/6J mice at baseline (Fig.4G). HFD treatment did not alter circulating levels of cytokines in any strain. The total antioxidant status (TAS) was measured in serum during daytime to inform about the capacity to neutralize free radicals associated to inflammation. In these conditions, basal levels of TAS were unexpectedly lower in WSB/EiJ than in C57BL/6J mice, whatever the feeding conditions (Fig.4H).

**Figure 4:**
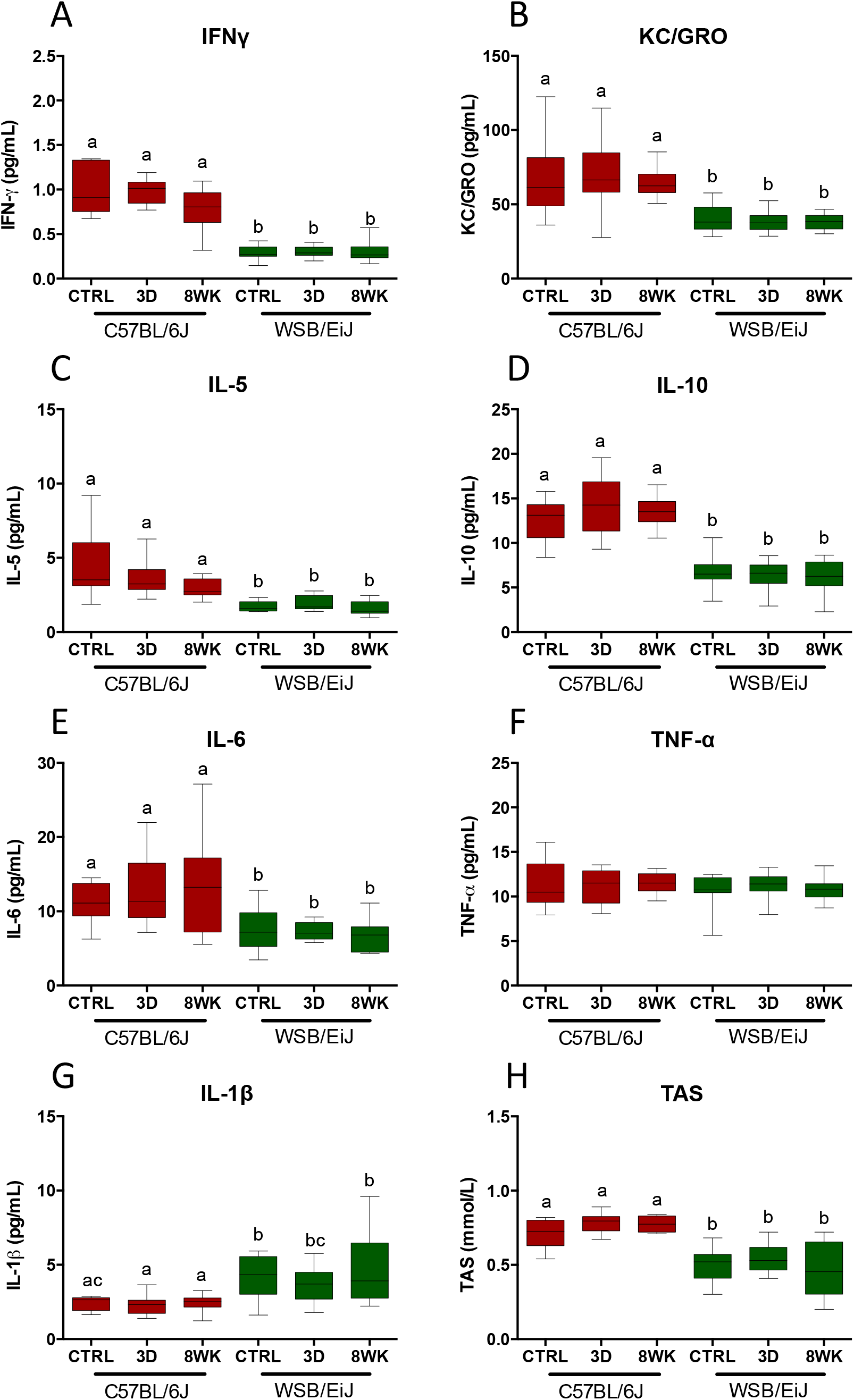
WSB/EiJ mice show overall lower circulating inflammatory markers as compared to C57BL/6J mice, with no effect of diet. Compared analysis of main circulating cytokines and inflammatory markers in twelve week-old C57BL/6J and WSB/EiJ male mice either maintained under control feeding (CTRL), or challenged with HFD for the last 3 days of the period (3D) or during the whole 8-week experiment duration (8WK). A. Interferon gamma (IFN-γ in pg/mL; N=11-13 per group); B. Keratinocyte-derived cytokine (KC/GRO in pg/mL; N=10-14 per group); C. Interleukin 5 (IL-5 in pg/mL; N=11-13 per group); D. Interleukin 10 (IL-10 in pg/mL; N=10-14 per group); E. Interleukin 6 (IL-6 in pg/mL; N=11-12 per group); F. Tumor Necrosis Factor alpha (TNF-α in pg/mL; N=11-14 per group); G. Interleukin 1 beta (IL-1β in pg/mL; N=11-14 per group). H. Total antioxidant status (TAS in mmol/L; N=10-12 per group). Graphs represent medians using box and whiskers. Significant differences were indicated by different letters to account for between strain and between diet group differences (P≤0.05; GLM).

Central indicators of inflammation were then compared among HFD mice from both strains. In agreement with peripheral inflammatory markers, C57BL/6J mice showed significantly higher levels of microglial activation than WSB/EiJ mice, an indicator of central inflammatory response. Indeed, analyses of IBA1 immunostaining revealed that microglia cell area and density in the arcuate nucleus (ARC) were higher overall in C57BL/6J compared to WSB/EiJ (Fig.5A-B). A 3d HFD challenge induced a slight but significant increase in microglia area, but not density, in both strains (Fig. 5B). The same effects of diet were observed in the paraventricular nucleus (PVN) for microglia area and density, but no strain effect was seen for the microglia area (Fig. 5C-D), whereas an interaction between strain and diet was significant for the microglia density in the PVN (P=0.005).

**Figure 5:**
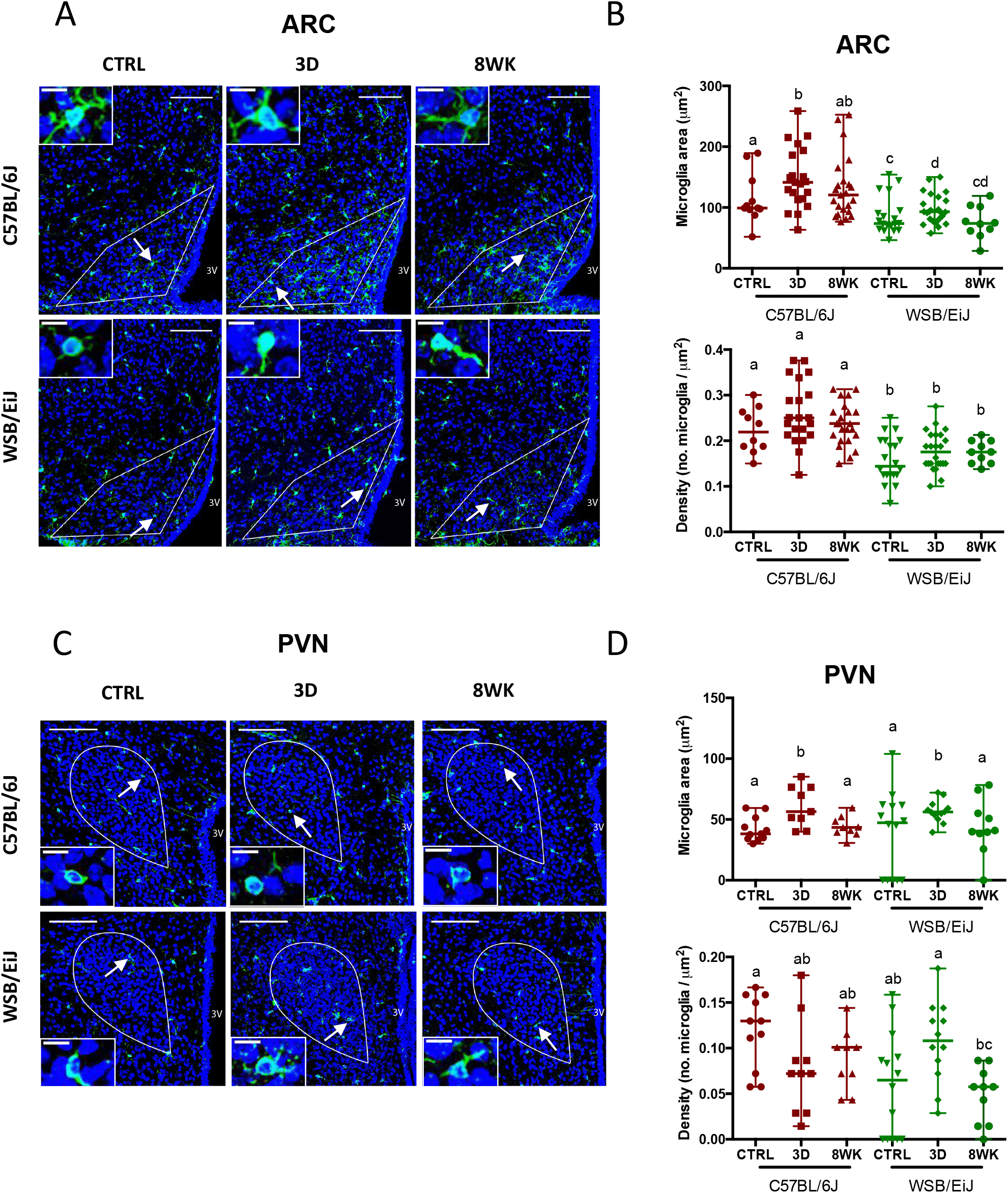
WSB/EiJ mice show no increase of inflammatory responses to HFD in the hypothalamic arcuate nucleus (ARC) in contrast with C57BL/6J mice, but comparable responses in the hypothalamic paraventricular nucleus (PVN). Immuno-histochemical analysis of microglia cells in the hypothalamic nuclei of twelve old C57BL/6J and WSB/EiJ male mice either maintained under control feeding (CTRL), or challenged with HFD for the last 3 days of the period (3D) or during the whole 8-week experiment duration (8WK). A. Representative confocal images of ARC immune-labelled with Iba1 (in green), a microglia marker, and DAPI-stained nuclei (in blue), in coronal sections of C57BL/6J (top panel) and WSB/EiJ (bottom panel) mice. Regions of interest (ROI) are delimited in white. Microglia pointed by arrowhead were magnified in insets at the top left of the main image; B. Quantitative analysis of microglia area (top; soma + ramifications, in μm2) and density (bottom; number of microglia per μm2 of ROI) obtained from confocal images showed in A (N = 10-24 sections/group). Graphs show median values with range in scatter dot plot. Significant differences are indicated by different letters to account for between strain and between diet group differences (P≤0.05; GLM); C. Representative confocal images of PVN immune-labelled with Iba1 (in green), a microglia marker, and DAPI-stained nuclei (in blue), in coronal sections of C57BL/6J (top panel) and WSB/EiJ (bottom panel) mice. Regions of interest (ROI) are delimited in white. Microglia pointed by arrowheads were magnified in insets at the bottom left of the main image; D. Quantitative analysis of microglia area (top; soma + ramifications, in μm2) and density (bottom; number of microglia per μm2 of ROI) obtained from confocal images showed in C (N = 10-24 sections/group). Graphs show median values with range in scatter dot plot. Significant differences are indicated by different letters to account for between strain and between diet group differences (P≤0.05; GLM); Scale bars in main images: 100μm; scale bars in insets = 10 μm.

### 3.3. Resistance to DIO in WSB/EiJ mice could be associated with enhanced transport and signalling of endocrine molecules in the hypothalamus

Microglia along with astrocytes and tanycytes provide support and protection for neighbouring neurons. GFAP labelling revealed a differential glial patterning between both strains in hypothalamic ARC (Fig.6A) and PVN (Fig.6B). Two types of cells were revealed: i) star-shaped cells corresponding to astrocytes (Fig.6, yellow arrows); ii) cells lining the 3^rd^ ventricle (3v) and further characterized by long radial projections in the parenchyma (Fig.6, white arrows). This latter type of cell could be tanycytes based on the fact that they covered the interface between the systemic compartment and hypothalamic neurons. Confocal microphotographic analysis revealed a sharply distinct morphology and distribution of these tanycyte-like cells between strains regardless of feeding conditions. WSB/EiJ mice displayed more profuse, linear and extended GFAP-immunoreactive cytoplasmic processes across the ARC and more particularly in the PVN, which may enhance transport of endocrine molecules. In contrast, C57BL/6J strain displayed a sparse labelling in ARC and especially in the PVN, with few and short GFAP-immunoreactive extensions, which stood close to the ventricular domain.

**Figure 6:**
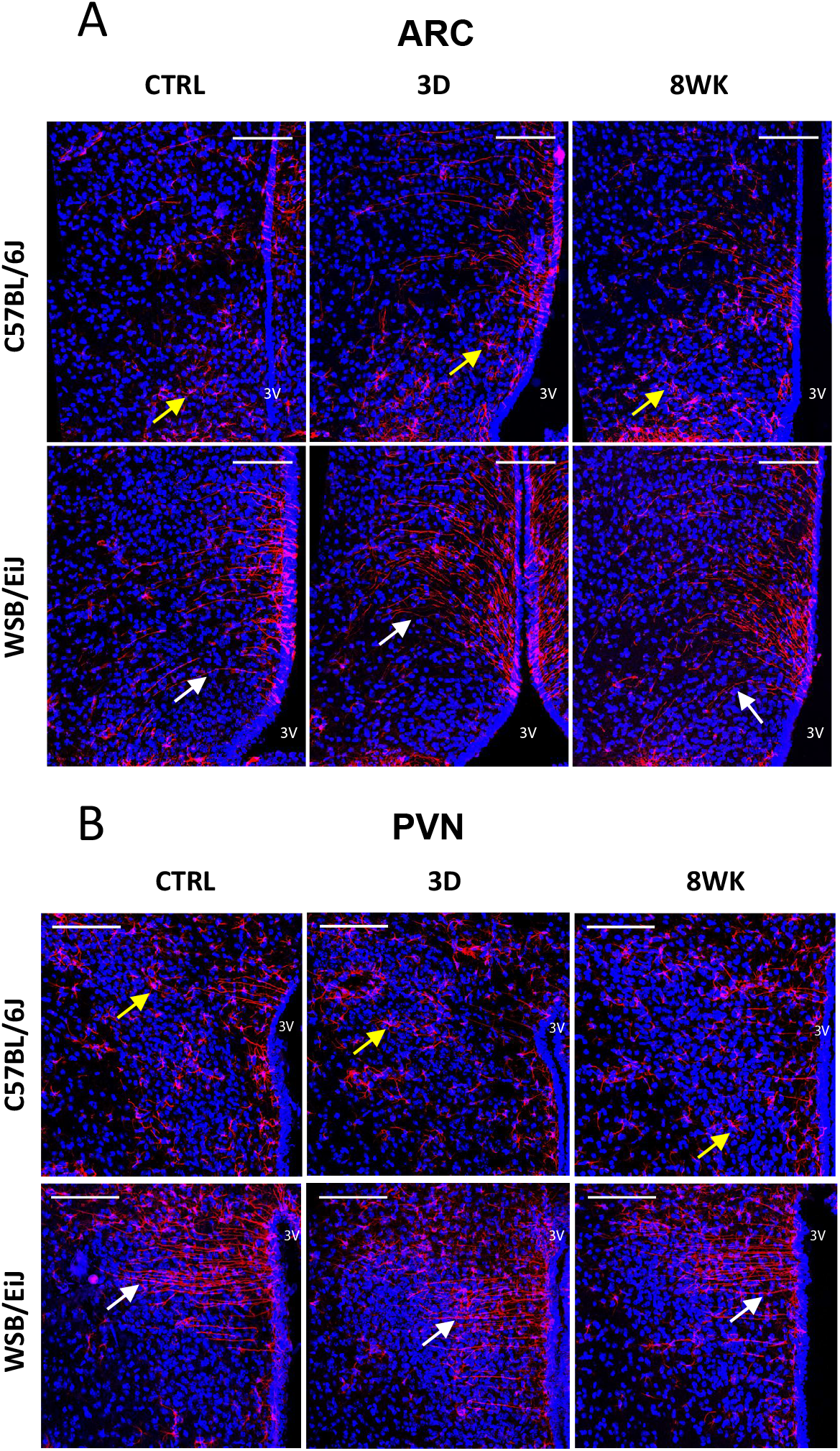
WSB/EiJ mice show remarkably well-defined elongated ependymal cells in both the hypothalamic arcuate nucleus (ARC) and the hypothalamic paraventricular nucleus (PVN), whatever the diet is. Immuno-histochemical analysis of GFAP-positive cells in the hypothalamic nuclei of twelve week-old C57BL/6J and WSB/EiJ male mice either maintained under control feeding (CTRL), or challenged with HFD for the last 3 days of the period (3D) or during the whole 8-week experiment duration (8WK). A. Representative confocal images of GFAP-positive cells (in red) and DAPI-stained nuclei (in blue) in the ARC region of coronal sections of C57BL/6J (top panel) and WSB/EiJ (bottom panel) mice. GFAP-positive elongated tanycyte-like ependymal cells (pointed by white arrowheads) were observable in WSB/EiJ sections in the top region of ARC close to the third ventricle (3v), contrasting with shorter and well distinguishable GFAP-positive astrocyte-like cells in C57BL/6J mice (pointed by yellow arrowheads). B. Representative confocal images of GFAP-positive cells (in red) and DAPI-stained nuclei (in blue) in the PVN region of coronal sections of C57BL/6J (top panel) and WSB/EiJ (bottom panel) mice. GFAP-positive elongated ependymal cells (pointed by white arrowheads) were observable in WSB/EiJ sections in the PVN region close to the third ventricle (3v), contrasting with shorter and well distinguishable GFAP-positive astrocyte-like cells in C57BL/6J mice (pointed by yellow arrowheads). Scale bars: 100μm.

### 3.4. WSB/EiJ hypothalamic mitochondria dynamics differ from C57BL/6J

Ultra-microscopic observations by transmission electron microscopy (TEM) revealed the presence of lipid droplets close to the 3v in the PVN of C57BL/6J mice under all diet conditions (Fig.S1A-B). In contrast, no or very few lipid droplets were observed in the same region in WSB/EiJ mice, being thus considered as negligible.

Mitochondrial architecture in the parvocellular PVN (Fig.S1C) was monitored by TEM and classified along four types as indicators of mitochondrial state of activity under obesogenic conditions [18, 19] (Fig.S1D). The two strains depicted a clear differential response to HFD in terms of mitochondria dynamics, revealed by a significant interaction between strain and diet regarding the ratio of orthodox versus condensed mitochondria (p<0.05). In the parvocellular region of the C57BL/6J PVN, the ratio of mitochondria types between condensed and orthodox remained stable after diet treatments. In contrast, the type of mitochondria surrounding the nuclear area of individual neuronal cell bodies varied and was diet-dependent in WSB/EiJ mice, indicating transiently enhanced proportion of active mitochondria (condensed) in parvocellular neurons of WSB/EiJ under obesogenic conditions: 3d HFD induced a significant transient increase in condensed type (p=0.001), at the expense of the orthodox category (p=0.006), but both returned to baseline levels with 8wk HFD. Regarding the global characteristics of mitochondria, neither aspect ratio (Fig.S1E) nor surface density coverage (Fig.S1F) showed any effect of diet or strain. Nevertheless, strain-independent-diet-induced changes in the area of mitochondria were seen (p<0.001, Fig.S1G), 8wk HFD inducing a decrease of the area of each individual mitochondrion compared to control diet. Finally, measuring the ATP content of ARC or PVN in both strains revealed that WSB/EiJ, but not C57BL/6J, increased their ATP content under 8wk HFD, in both ARC and PVN (Fig.S1H-H’).

### 3.5. Hypothalamic inflammation and mitochondrial gene pathways are differentially regulated by HFD in WSB/EiJ and C57BL/6J

To study whether the observed phenotypic characteristics of the WSB/EiJ could be explained by differential regulatory pathways, we analysed the expression of 86 genes related to inflammation and mitochondria in both strains under HFD, focusing on the key hypothalamic nuclei involved in central control of metabolism, the PVN and ARC.

PCA analysis of microfluidic qRT-PCR data for PVN and ARC datasets was done and dCT values plotted on two-dimensional axis (Fig.S2). PCA analysis of the ARC or the PVN datasets clearly discriminated between strains (with 37 % and 31 % of the variability explained by the two dimensions, respectively). This result confirmed the strain specificity of the gene expression profiles in the ARC and PVN, with a strain differential response to diet attested by an extremely significant interaction between both factors (strain and diet, ARC: p = 8E^-4^; PVN : p = 3E^-6^). Interestingly, the analysis of the list of genes contributing to the weights of PCA dimensions allowing for the strains discrimination revealed a clear differential expression pattern between the ARC and PVN (Tables S1-2). Most of these genes were significantly differentially expressed as a function of strain, and some even showed a significant interaction between strain and diet (p< 0.05, Table S1-2, 3^rd^ column, and Fig.S3-6 for graphs, see below for detail on gene expression).

We next analysed the expression data so as to determine the pathways implicated in the differential gene expression profiles between strains. For both the ARC and the PVN, heatmaps showed distinct differential expression profiles as a function of strain and diet. Interestingly, most regulated pathways identified were not only related to mitochondria and inflammation, as expected regarding the gene selection we made, but also to metabolism. For the ARC, the significantly regulated pathways (see Fig.7A for p values), beside those related to inflammation (adipocytokine pathway, cytokine-cytokine receptor pathway, Hippo signalling pathway) and mitochondria (oxidative phosphorylation) were the metabolic, FoxO, MAPK and AMPK signalling pathways. When looking more closely to the genes involved in the AMPK pathway (Fig.7B), we observed a significant effect of strain for three (*Irs 1, mTor* and *Insr*) out of four genes, and even a significant interaction between the strain and the diet for *Insr* (being more expressed but not regulated by HFD in the WSB/EiJ, whereas in C57BL/6J, it was increased by 8wk HFD to the level of WSB/EiJ). In the case of the PVN (Fig.8A for p values), certain signalling pathways apart from inflammation (adipocytokine pathway) and mitochondrial (oxidative phosphorylation) pathways displayed regulations notably AMPc, Jak-STAT, PI3K-AKT, FoxO and metabolic pathways. Of the five genes involved in the FoxO pathway (mediating cellular processes as apoptosis, glucose metabolism, oxidative stress resistance and longevity), and showing a similar pattern of expression (more expressed in WSB/EiJ CTRL and even more in 3d HFD) three (*Tgfb1, Foxo3* and *Prkaa1*) were significantly differentially expressed between the groups (Fig.8B). For *Tgfb1* expression, a strong interaction (p=0.004) between diet and strain was observed. *Tgfb* expression was not affected by HFD in C57BL/6J, whereas in WSB/EiJ CTRL and 3d HFD, *Tgfb* was expressed two-fold more than in C57BL/6J, and was repressed to the level of C57BL/6J by 8wk HFD. *Foxo3* levels were significantly higher in WSB/EiJ, but not regulated by HFD, whereas *Prkaa1* expression showed also a significant interaction between the strain and the diet, particularly for the 3d HFD groups, where *Prkaa1* was 1.5 times more expressed in WSB/EiJ than in C57BL/6J (Fig.8B). Regarding the oxidative phosphorylation and metabolic pathways, a remarkable expression pattern is the strong increase denoted by the red colour in the 8wk HFD WSB/EiJ (Fig.8A,C).

**Figure 7:**
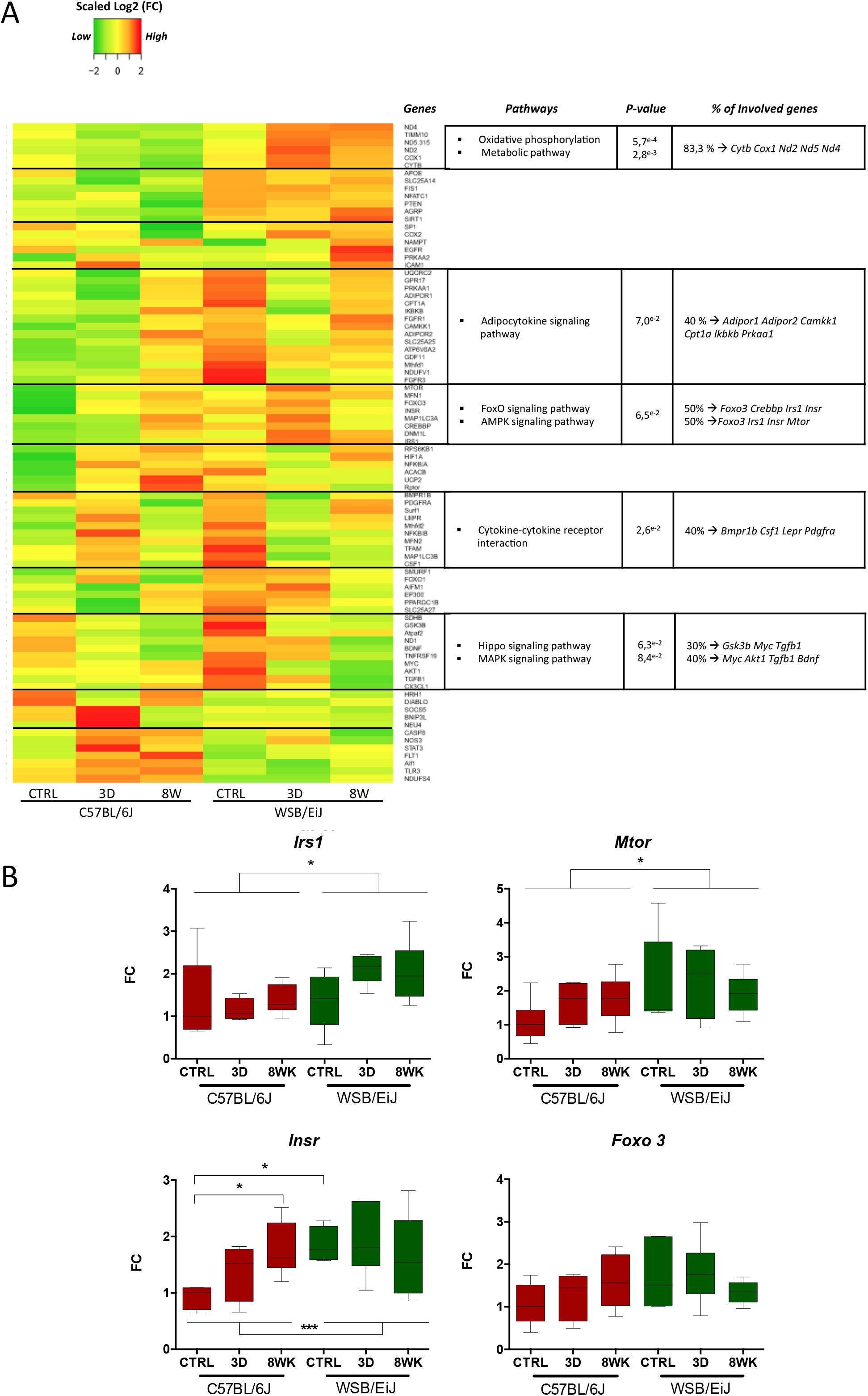
Expression profiles of genes related to inflammation or mitochondria are different between both strains, and also differentially influenced by HFD in the ARC. ***A. Heatmap showing for the arcuate nucleus of the hypothalamus (ARC), the relative gene expression data (expressed as scaled log2 Fold Change values) for 86 genes related to inflammatory and mitochondrial pathways:*** Data were collected by microfluidic qRT-PCR from ARC of twelve week-old C57BL/6J (left panel) and WSB/EiJ (right panel) mice either maintained under control feeding (CTRL), or challenged with HFD for the last 3 days of the period (3D) or during the whole 8-week experiment duration (8WK). Color transition represents gene expression levels expressed as scaled log2 Fold Change values (scaled Log2(FC)): green for low expression, yellow for mean expression and red for high expression. Genes with similar expression profiles were clustered together and the cluster delimited manually by a black horizontal line. Each cluster of genes was analysed against the 86 genes with DAVID bioinformatics tools and significant pathways were identified. The summary of the significant pathways identified is shown in the table (at the right of each cluster) with the associated significant P-value (P≤0.1), the percentage of genes in the cluster involved in each pathway, and the names of the involved genes. ***B. Individual gene expression data for representative significantly differentially regulated genes involved in pathways defined in (A):*** Data were collected by microfluidic qRT-PCR from microdissected ARC of twelve week-old C57BL/6J (red) and WSB/EiJ (green) mice either maintained under control feeding (CTRL), or challenged with HFD for the last 3 days of the period (3D) or during the whole 8-week experiment duration (8WK). *Irs1:* Insulin receptor substrate 1; *Mtor*: mammalian target of rapamycin; *Insr*: insulin receptor; *Foxo3*: Forkhead box protein O3. Two-way Anova analysis revealed a significant strain effect (independently of the diet) was observed for *Mtor* and *Irs1. Insr* expression was influenced both by the strain and the diet, indicating a strain difference in the response to the diet, with no effect of the diet in WSB/EiJ, whereas HFD induced a high increase in *Insr* expression in C57BL/6J. Post Hoc tests results are indicated on the graph. (*, P≤0.05; ***, P≤0.001. N=5-7 per group), FC : relative fold-change expression.

**Figure 8:**
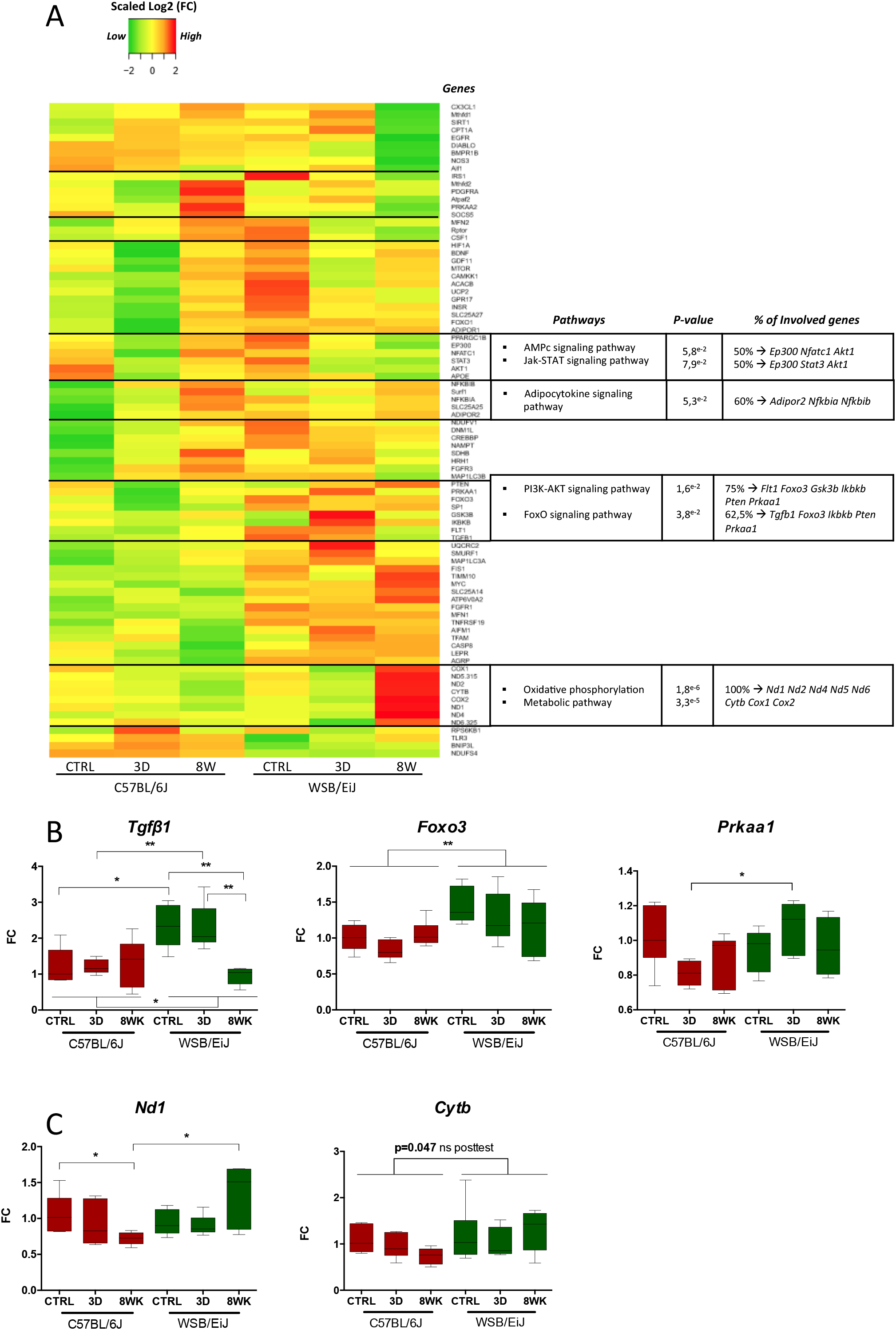
Expression profiles of genes related to inflammation or mitochondria are different between both strains, and also differentially influenced by HFD in the PVN. ***A. Heatmap showing for the paraventricular nucleus of the hypothalamus (PVN), the relative gene expression data (expressed as scaled log2 Fold Change values) for 86 genes related to inflammatory and mitochondrial pathways:*** Data were collected by microfluidic qRT-PCR from PVN of twelve week-old C57BL/6J (left panel) and WSB/EiJ (right panel) mice either maintained under control feeding (CTRL), or challenged with HFD for the last 3 days of the period (3D) or during the whole 8-week experiment duration (8WK). Color transition represents gene expression levels expressed as scaled log2 Fold Change values (scaled Log2(FC)): green for low expression, yellow for mean expression and red for high expression. Genes with similar expression profiles were clustered together and the cluster delimited manually by a black horizontal line. Each cluster of genes was analysed against the 86 genes with DAVID bioinformatics tools and significant pathways were identified. The summary of the significant pathways identified is shown in the table (at the right of each cluster) with the associated significant P-value (P≤0.1), the percentage of genes in the cluster involved in each pathway, and the names of the involved genes. ***B. Individual gene expression data for representative significantly differentially regulated genes involved in pathways defined in (A):*** Data were collected by microfluidic qRT-PCR from microdissected PVN of twelve week-old C57BL/6J (red) and WSB/EiJ (green) mice either maintained under control feeding (CTRL), or challenged with HFD for the last 3 days of the period (3D) or during the whole 8-week experiment duration (8WK). *Tgfβ*: Transforming growth factor beta Transcription factor; *Foxo3:* forkhead box O-3; *Prkka1:* Protein kinase AMP-activated catalytic subunit alpha 1; *Nd1:* NADH-ubiquinone oxidoreductase chain 1; *Cytb*: Cytochrome b. Two-way Anova analysis revealed a significant combined effect of strain and diet on *Tgfβ* and *Nd1* expressions, whereas *Foxo3* expression is significantly different in the two strains independently of the diet, and *Prkka1* expression is regulated by the diet, independently of the strain. Post Hoc tests results are indicated on the graph. (*, P≤0.05; **, P≤0.01. N=5-7 per group). FC : relative fold-change expression.

The heatmaps also revealed that several genes followed similar patterns of regulation, even if no significant pathway was identified. In the ARC (Fig.7A), some genes showed higher expression in WSB/EiJ, without regulation by HFD (detailed in Fig.S3). These genes were associated in our initial GO term analysis either to mitochondria (Fig.S3A, notably *Slc25a14* (coding for Ucp5) *Fis1* and *Timm10*, involved in mitochondrial activity, but also *Sirt1*, which is protective against weight gain), or to inflammation (Fig.S3B, notably *Apoe*, involved in catabolism of lipoprotein particles and lipid transport and *Agrp*, an orexigenic factor). In contrast, some other genes also related to mitochondria or inflammation were significantly less expressed in WSB/EiJ and not regulated by HFD (Fig.S5A-B notably *Diablo*, implicated in apoptotic pathways and *Aif1*, gene coding for Iba1, marker of activated microglia). Some genes, as *Bdnf, Egfr* and *Fgfr3* (factors involved in proliferation) were also differentially regulated in response to HFD (Fig.S6). The regulation patterns seen in the ARC were also observed in the PVN (Fig.8A), with several genes related to mitochondria or inflammation being more (Fig.S4A, as *Slc25a14, Fis1* and *Timm10*, as already seen in the ARC, and *Agrp, AdipoR1, LepR* and *InsR*, all four main actors of metabolic homeostasis) or less (Fig.S5C-D, notably factors involved in apoptosis or inflammatory pathways, as *Diablo, Tlr3* and *Bmpr1b*) expressed in the WSB/EiJ. Interestingly, more genes were significantly differentially regulated by HFD in the PVN than in the ARC (Fig.S6), as *Ppargc1b* (associated with resistance to obesity) and *Nampt* (which is known to activate insulin receptor and has insulin-mimetic effects, lowering blood glucose and improving insulin sensitivity). Interestingly, these two genes showed higher levels in PVN of WSB/EiJ control and were decreased significantly (or not) by 8wk HFD, whereas 8wk HFD induced an increase in C57BL/6J to reach an expression level close to WSB/EiJ control levels.

## 4. DISCUSSION

We compared an obesity-resistant strain of mice (WSB/EiJ) with an obesity-prone C57BL/6J strain, exploiting the differences between strains in response to an HFD challenge, with the aim of identifying the regulatory mechanisms accounting for differences in susceptibility to DIO. Firstly, we verified that, in our laboratory conditions, we could reproduce the DIO resistance of the WSB/EiJ mice, observed by Lee et al [12]. Three days under HFD were sufficient to induce body mass increase in C57BL/6J mice, mostly due to fat accumulation in eWAT, an early sign of obesity. In contrast, the results confirmed the DIO-resistant phenotype of WSB/EiJ mice [12], as the challenge failed to induce WAT accumulation and concurrent gain of body mass even after eight weeks of HFD. Our results go far beyond those of Lee et al. in that we show not only differences in fat accumulation, but also reduced circulating lipids and leptin, a lower inflammatory status and an increased mitochondrial activity in the WSB/EiJ compared to the C57BL/6J mice.

A first significant finding was the absence of accumulation of ectopic fat in WSB/EiJ mice under obesogenic conditions. This feature was associated with specific lipid turnover and correlated with lower circulating lipids and leptin, but increased hydroxybutyrate levels, as compared to C57BL/6J. As hydroxybutyrate is a marker of fatty acid oxidation [20] and is usually used as a metabolic substrate in periods of starvation, our results support the hypothesis that WSB/EiJ mice would be constantly under a fasting phenotype. This is further supported by the lower levels of anorexigenic signals (leptin, PYY and GIP) in WSB/EiJ, as well as the higher mRNA expression of the orexigenic *Agrp* in the ARC of WSB/EiJ mice (Fig.S3B).

Among the endocrine parameters evaluated, leptin could play a pivotal role in the differential response between strains during the initial phase of HFD treatments. As expected, regarding the increase in the percentage of eWAT, leptin levels were increased in C57BL/6J mice upon 3 days and 8 weeks of HFD, contrasting with steady low and unchanged levels in WSB/EiJ mice. Sustained high leptin levels, in cross-talk with insulin resistance, have previously been related to obesity progression [21]. The leptin response to HFD in C57BL/6J, as a consequence of expanded adipocytes, failed to efficiently induce the expected anorexigenic effects, suggesting the early settlement of central leptin resistance in C57BL/6J mice. In the same line, differential response in peripheral glucose metabolism has previously been observed between C57BL/6J and WSB/EiJ mice. This difference, confirmed in our study, was more pronounced after a longer exposure to HFD (16 weeks of HFD) which allowed the establishment of insulin resistance in C57BL/6J strain, accompanied by increased total adiponectin levels [22, 23]. Our results further revealed a significant inverted pattern between strains for circulating C-peptide 2 levels, which is secreted in equimolar concentrations to insulin [24]. Adiponectin is known to accelerate clearance of HFD-derived fatty acid plasmatic overload [22]. The observed protective increase in adiponectin was transient in both strains, as assessed by a return to normal baseline levels upon 8wk treatment. This suggests that this short-lived peripheral hormone may be an immediate early response cue to obesogenic treatment. The differences between strains previously reported could appear after sustained long term response to HFD [22].

The role of resistin in obesity and adipogenesis is controversial. As an adipose tissue-derived hormone, one might expect an increase of resistin levels in C57BL/6J mice after 8 weeks under HFD [25]. However, despite initial increase after 3 days HFD, no significant variation was observed in this hormone over longer periods in C57BL/6J mice. No difference was observed in WSB/EiJ under any treatment, but it is of note that the levels at 3d were lower in the WSB/EiJ compared to the C57BL/6J, which is in accordance with the changes in leptin levels.

Our observations suggest that WSB/EiJ mice may benefit from a basal phenotype promoting the use of lipids, therefore triggering a fast and efficient response to a load of circulating lipids. Such phenotype could originate at the central level, and more particularly in hypothalamic regions involved in lipid metabolism and energy homeostasis. Our gene expression analysis suggests an increased hypothalamic sensitivity for WSB/EiJ mice to homeostatic signals, such as leptin, insulin and adiponectin, the latter promoting decreased body weight by acting on the PVN [26]. Indeed, their receptors (respectively *LepR, InsR* and *AdipoR1*) were more expressed in the PVN, and also in the ARC (for *InsR*) of WSB/EiJ mice as compared to C57BL/6J. Genes involved in the insulin signalling pathway (*InsR, Irs1, mTor, Nampt*) were regulated in the same way, being more expressed in the hypothalamus of WSB/EiJ mice, again suggesting an increased sensitivity to insulin signalling, already at baseline [27]. Overall, gene expression analysis suggests an increased hypothalamic sensitivity to peripheral hormonal and nutrient signals in WSB/EiJ mice, which could in part explain their resistance to diet induced obesity. This increased sensitivity should be sustained by a more efficient nutrient and hormonal transport network, allowing for faster sensing of variations in metabolic conditions than C57BL/6J mice. Tanycytes, specialized ependymal cells lining the wall of the third ventricle of the hypothalamus, play a major role in the control of food intake and energy expenditure, mainly involved in hypothalamic nutrient sensing and shuttling of metabolic signals from periphery to hypothalamic neurons [28]. Interestingly, we observed very well elongated tanycytes with cytoplasmic extensions in the hypothalamus of the WSB/EiJ strain. PVN neurons of WSB/EiJ mice were profusely covered with cytoplasmic extensions spanning from the paraventricular territory, which were not observed in C57BL/6J mice, regardless of metabolic conditions. ARC also displayed such differential labelling in both strains but at a lesser extent that in PVN, probably because the primary type of tanycytes in this region are β-tanycytes rather than α-tanycytes [29]. This result supports the hypothesis for increased hypothalamic sensitivity and an enhanced transport of endocrine molecules in WSB/EiJ in comparison to C57BL/6J.

Moreover, one clear consequence of the obesogenic treatment in C57BL/6J was the rapid accumulation of lipid droplets along the region bordering the 3v, which connects the cerebrospinal fluid to neuroendocrine systems. In contrast, WSB/EiJ did not accumulate significant amounts of droplets in this hypothalamic region as a consequence of any diet. Hypothalamic lipid accumulation can be detected by nutrient-sensitive glial cells and neurons, promoting inflammatory responses [30] and further dampening endocrine signalling [31–33]. The absence of lipid droplets in WSB/EiJ is consistent with either a peripheral or a hypothalamic enhanced lipid metabolism, therefore preventing from peripheral and central accumulation of fat. An interesting result in line with this hypothesis for a contribution of hypothalamic lipid metabolism in the WSB/EiJ phenotype is the higher *Apoe* expression in the ARC of WSB/EiJ compared with C57BL/6J. As *Apoe* plays a major role in lipid metabolism and transport from the astrocytes to the neurons [34], we cannot rule out that the efficiency of central lipid metabolism and transport could contribute to the WSB/EiJ DIO resistance.

HFD is known to induce low-grade inflammatory responses (called meta-inflammation) in immune and metabolic cells from several tissues including adipose tissue, liver, muscle, pancreas, gastrointestinal tract and also in the brain, and become chronic concomitant with the settlement of obesity [35, 36]. Under our obesogenic conditions we observed low circulating levels of cytokines in WSB/EiJ mice, consistent with the lack of immune and inflammatory responses, and lack of obesity. On the contrary, C57BL/6J displayed higher levels of most of the tested circulating cytokines including IFN-γ, KC/GRO, IL-5, IL-10 and IL-6. This proinflammatory state might contribute to explain increased leptin levels as soon as HFD was provided and therefore concur for the higher susceptibility to obesity and insulin resistance in this strain [35]. IL-1β was the only analysed cytokine displaying higher levels in WSB/EiJ mice compared to C57BL/6J. This effect could be explained by the fact that IL-1β is a pleiotropic cytokine involved in the inflammatory response, metabolic regulation and exerts anorexigenic and pyrogenic roles consistent with the WSB/EiJ lean phenotype [37].

Despite the higher levels observed in C57BL/6J, none of the two strains displayed significant changes in circulating inflammatory cytokines under acute obesogenic conditions, as opposed to that previously reported by others [38, 39], even if leptin and %WAT were increased. Several elements may account for this discrepancy. First, a highly intense fat overload of 60% was provided in those studies, instead of 45% tested here. Second, the control food used in our study was low fat content but the same high carbohydrate content than the HFD food (17% Kcal from carbohydrates). As sugar consumption is known to induce inflammation [40], a high sucrose-low fat diet could settle, under control conditions, a higher basal inflammatory status in C57BL/6 mice, potentially buffering the effect of fat on inflammatory markers.

Third, a putatively significant increase in the immediate-early cytokine response may be transient and subjected to high variations in blood samples during the initial stages of low-grade meta-inflammation. In harmony with our findings, Lee et al reported that such inflammatory peripheral signalling was most important during the chronic phase to mediate metabolic insulin resistance [39]. Indeed, Thaler et al showed previously that hypothalamic low-grade inflammation (e.g. in ARC/PVN microglia) concomitantly to what we observed, occurs as a rapid response to HFD but requires a certain length of time and intensity to be reflected at a peripheral level [7].

The central coordination of energy homeostasis involves ARC and PVN hypothalamic nuclei, as well as neuro-hormonal synchronizing signals, which adjust the responses centrally and peripherally. Hypothalamic neurons depend upon proper interaction with their surrounding glia for both support and paracrine aspects [32]. Microglial cells have been revealed as main components linking metabolism and inflammation under obesogenic conditions. In addition to a role in lipid sensing, microglia also participate in the inflammatory process, acting as the macrophage-like immune cells of the brain. Activated microglia proliferate, change their morphology and release cytokines in response to immune threats, such as lipid overload [7, 41, 42]. In response to HFD, activated microglia orchestrate the regulation of food intake, modifying leptin signalling and neural function [32]. As short as 1 to 3 days HFD have been reported to activate microglia in the ARC of rats [7]. However, to our knowledge, no parallel information is available for the PVN. Here, we quantified microglia density and cellular area in ARC and PVN after an obesogenic challenge in both strains. Concomitantly with circulating inflammatory markers, C57BL/6J displayed signs of enhanced activation of microglia in ARC under control conditions, which was correlated with a higher *Aif1* mRNA expression in the ARC of C57BL/6J mice. Such chronic, low-grade inflammation both within the hypothalamus and in periphery, may contribute to the C57BL/6J strain’s inherent predisposition to insulin resistance and the associated leptin-impaired response to HFD [3]. Moreover, HFD treatment of C57BL/6J mice elicited even higher microglial activation in the ARC, whereas WSB/EiJ exhibited a modest but significant increase in microglial activation, but only after 3d HFD, not at 8wk. This increase could act as a trigger for a defensive response to HFD, allowing a rapid corrective action to adjust to the fat overload. In C57BL/6J, this increase was maintained after 8wk HFD, consistent with an increased inflammatory state associated with DIO in this mouse strain.

A rapid response to 3d HFD was also observed in mitochondria of the PVN parvocellular neurons in WSB/EiJ mice, supporting the idea of an enhanced signalling in hypothalamic command centres allowing adapted neuronal responses to HFD. Mitochondria are key dynamic organelles involved in energy supply at sites of high ATP consumption, associated with neuronal activity and involved in mediating hypothalamic responses to food intake and metabolic state [10]. The morphology of mitochondria in these zones changed to a more active state in WSB/EiJ mice in response to HFD, indicating an increased flexibility in activity that may be associated with neuronal responses to energy demand [7, 32]. The more active state of the WSB/EiJ hypothalamic mitochondria was confirmed by an increased ATP production after 8wk HFD and the mitochondrial gene expression analysis. Indeed, most of the genes involved in mitochondrial activity were increased in the WSB/EiJ (*Slc25a14, Fis1, Timm10, Sirt1*, …), both in ARC and PVN, and even more increased after 8wk HFD. Of particular interest, is the large increase in genes involved in the FoxO signalling and oxidative phosphorylation pathways observed in WSB/EiJ 3d and/or 8wk HFD, in both hypothalamic nuclei. This can be interpreted as an adaptive response to nutrient excess, leading to changes in mitochondrial metabolism.

Overall, the gene expression results emphasise that these two strains are characterized by extremely different hypothalamic transcriptomic profiles, not only for the genes involved in inflammation and mitochondrial activity, but also for the genes involved in metabolic regulation. As hypothesized, we found that the mechanisms involved in metabolic homeostasis could be different between the strains, but also that the HFD challenge induced differential regulation between the strains with regards to the pathways related to metabolism.

## 5. CONCLUSIONS

These observations lead to the conclusion that enhanced hypothalamic mitochondrial activity, associated with efficient transport and signalling of endocrine molecules, could act as a protective response against fat overload and the settlement of obesity-associated low-grade meta-inflammation.

## Acknowledgements

Authors wish to thank the valuable technical support from the following French scientific services: Cytometry and Immunobiology Facility (Cybio, INSERM U1016, Paris) and the Plateforme de Biochimie (CRI, INSERM UMR1149, Paris). We also acknowledge the ImagoSeine Facility, (member of the France BioImaging infrastructure supported by the French National Research Agency, ANR-10-INSB-04, “Investments for the future”). The expert animal care provided by S. Sosinski and F. Uridat is gratefully acknowledged. This work was supported by European Union contracts Nina (GA-238665), Switchbox (GA-259772) and Human (GA-602757). L. Chamas received a PhD Fellowship from Ile de France Region. The authors declare there is no conflict of interest that would prejudice the impartiality of this scientific work.

## Abbreviations

HFD: high fat diet;
CTRL: control diet;
DIO: diet-induced obesity;
ARC: Arcuate nucleus of the hypothalamus;
PVN: paraventricular nucleus of the hypothalamus;
eWAT: ependymal white adipose tissue;
iWAT: inguinal white adipose tissue;

