## Supplementary Material for "Specific inflammatory and mitochondrial signatures characterise the WSB/EiJ mice diet-induced obesity resistance capacity"

### 1. Material and Methods

##### 1.1. Electron Microscopy Analysis

PVN regions were dissected under binocular and samples were prepared according to Combès et al., 2012, fixed overnight in 2.5% glutaraldehyde solution in Sörensen buffer, and post-fixed in 1% osmium tetroxide. After serial ethanol dehydration, PVN samples were embedded in epoxy mixture (Spurr's resin). Ultrathin sections were contrasted with uranyl acetate and observed with a transmission electron microscope (Hitachi H-7100). Pictures were taken using a Hamamatsu CCD camera, focusing in two zones of the PVN: the tanycyte region lining the third ventricle and the parvocellular region. In both regions of interest, a series of 10-12 photos were analysed per animal using Image J software, and two animals were used for each strain/treatment. Two independent observers blindly counted the presence (number and size) of lipid droplets in the tanycyte region. Mitochondrial type was assessed in the parvocellular neurons of the PVN. Mitochondria were classified as orthodox, condensed, on instances of fusion/fission or autophagy (mitophagy) events. In individual parvocellular neuronal cell bodies, images were taken around the nucleus to assess in the surrounding mitochondria: aspect ratio (AR, major axis/minor axis), coverage (total area of mitochondria/cytosolic area) and mean area of individual mitochondria were calculated according to Dietrich et al., 2013.

##### 1.2. ATP measurements

ATP content in hypothalamic tissues was measured using ENLITEN® ATP Assay System Bioluminescence Detection Kit (Promega, United States) following supplier's protocol. Briefly, frozen manually dissected PVN and ARC regions were lysed with a TissueLyser (Qiagen) in 400 μL of Reporter Lysis Buffer (Promega), using stainless steel beads, for 2 min at a 30 Hz frequency. Lysates were centrifuged 10 min at 10000g, at 4°C, and ATP measured in 10 μL of supernatant added by 100 μL of reagent using a luminometer (Berthold), and ATP concentration was obtained using a standard curve. Protein concentration was measured in each sample, using Pierce™ BCA Protein Assay Kit (Thermo Scientific), in order to normalize results.

1.3. **Bioinformatic data analysis :** *PCA:* A custom R tool was constructed to perform the principal component analysis (PCA) using R packages: the FactoMineR (version 1.35) and missMDA (version 1.11). dCT values of inflammation and mitochondria genes were plotted on two-dimensional axis (Dim 1, Dim 2).

#### 2. Supplementary figures legends:

**Figure S1. Mitochondria are more responsive to HFD in the WSB/EiJ strain than in the C57BL/6J in PVN region**. Compared analysis of mitochondrial parameters in twelve week-old C57BL/6J and WSB/EiJ male mice either maintained under control feeding (CTRL), or challenged with HFD for the last 3 days of the period (3D) or during the whole 8-week experiment duration (8WK). A-G. TEM analysis of mitochondria in parvocellular neurons of the PVN: A. Representative TEM image of presence (left panel) or absence (right panel) of lipid droplets (indicated with red arrows) in C57BL/6J and WSB/EiJ mice, respectively; B. Quantitation of the number of lipid droplets revealing no lipid droplet accumulation in the WSB/EiJ strain as opposed to C57BL/6J mice C. Representative TEM image of fusion-like events observed in WSB/EiJ control-fed mice; D. Quantitation of mitochondrial states in parvocellular neurons of the PVN. Mitochondria in WSB/EiJ were more responsive to HFD, and transiently changed morphology to more active states: in WSB/EiJ lower percentage of condensed mitochondria (found under conditions of high activity) transiently increased after 3d HFD to C57BL/6J levels, at expenses of percentage of orthodox states (normally found under standard conditions and activity); E. Aspect Ratio of mitochondria calculated as the ratio between mitochondria major axis and minor axis; F. Mitochondria surface density or coverage calculated as the ratio between area covered by mitochondria and the total cytosolic area; G. Area of individual mitochondria. No significant difference in mitochondrial aspect ratio or coverage were detected. However, HFD decreased size of mitochondria in both strains, suggesting fission induced by HFD exposure. H-H’. Analysis of the ATP content of

ARC (H) or PVN (H') in both strains revealed that WSB/EiJ, but not C57BL/6J, increased their ATP content under 8 weeks HFD. Significant differences were indicated by different letters on top of bars (P≤0.05).

**Figure S2. Principal component analysis (PCA) discriminates WSB/EiJ and C57BL/6J strains on the basis of hypothalamic gene expression**. Genes representative of inflammation and mitochondrial pathways were computed in a PCA analysis to verify whether the response to HFD differed between WSB/EiJ and C57BL/6J mice in ARC (upper panel) and PVN (lower panel). The individual factor maps revealed a clear discrimination between both strains around Dimension 2, especially in PVN. However, the response to HFD could not really be identified on the basis of PCA analysis. The contribution of each gene along PCA dimensions 1 and 2 is represented in Table S1 for ARC and Table S2 for PVN.

**Figure S3. Individual gene expression data for genes significantly more expressed in the ARC of WSB/EiJ mice compared to C57BL/6J mice**: Data were collected by microfluidic qRT-PCR from microdissected ARC of twelve week-old C57BL/6J (red) and WSB/EiJ (green) mice either maintained under control feeding (CTRL), or challenged with HFD for the last 3 days of the period (3D) or during the whole 8-week experiment duration (8WK). A: genes more expressed in the WSB/EiJ, related to mitochondria. B: genes more expressed in the WSB/EiJ, related to inflammation. Gene expression data were analyzed using two-way Anova to test the effects of diet, strain and their interaction. Post Hoc tests results are indicated on the graph (*, P≤0.05; **, P≤0.01; ***, P≤0.001. n=5-7); FC : relative fold-change expression.

**Figure S4. Individual gene expression data for genes significantly more expressed in the PVN of WSB/EiJ mice compared to C57BL/6J mice**: Data were collected by microfluidic qRT-PCR from microdissected PVN of twelve week-old C57BL/6J (red) and WSB/EiJ (green) mice either maintained under control feeding (CTRL), or challenged with HFD for the last 3 days of the period (3D) or during the whole 8-week experiment duration (8WK). A: genes more expressed in the WSB/EiJ, related to mitochondria. B: genes more expressed in the WSB/EiJ, related to inflammation. Gene expression data were analyzed using two-way Anova to test the effects of diet, strain and their interaction. Post Hoc tests results are indicated on the graph (*, P≤0.05; **, P≤0.01; ***, P≤0.001. n=5-7); FC : relative fold-change expression.

**Figure S5. Individual gene expression data for genes significantly less expressed in the ARC and PVN of WSB/EiJ mice compared to C57BL/6J mice** Data were collected by microfluidic qRT-PCR from microdissected ARC and PVN of twelve week-old C57BL/6J (red) and WSB/EiJ (green) mice either maintained under control feeding (CTRL), or challenged with HFD for the last 3 days of the period (3D) or during the whole 8-week experiment duration (8WK). A: genes less expressed in the ARC of WSB/EiJ, related to mitochondria. B: genes less expressed in the ARC of WSB/EiJ, related to inflammation. C : genes less expressed in the PVN of WSB/EiJ, related to mitochondria. D : genes less expressed in the PVN of WSB/EiJ, related to inflammation. Gene expression data were analyzed using two-way Anova to test the effects of diet, strain and their interaction. Post Hoc tests results are indicated on the graph (*, $P \leq 0.05$; **, $P \leq 0.01$; ***, $P \leq 0.001$. n=5-7); FC : relative fold-change expression.

**Figure S6. Individual gene expression data for genes significantly differentially regulated by HFD in the ARC and PVN of WSB/EiJ mice compared to C57BL/6J mice:** Data were collected by microfluidic qRT-PCR from microdissected ARC and PVN of twelve week-old C57BL/6J (red) and WSB/EiJ (green) mice either maintained under control feeding (CTRL), or challenged with HFD for the last 3 days of the period (3D) or during the whole 8-week experiment duration (8WK). A: genes differentially regulated in the ARC of WSB/EiJ, compared to C57BL/6J. B: genes differentially regulated by HFD in the PVN of WSB/EiJ compared to C57BL/6J, related to Mitochondria C : genes differentially regulated by HFD in the PVN of WSB/EiJ compared to C57BL/6J, related to Inflammation. Gene expression data were analyzed using two-way Anova to test the effects of diet, strain and their interaction. Post Hoc tests results are indicated on the graph (*, $P \leq 0.05$; **, $P \leq 0.01$; ***, $P \leq 0.001$. n=5-7); FC : relative fold-change expression.

#### 3. Supplementary tables legends

**Table S1**. Summary table of the statistics of the genes involved in inflammation and mitochondrial pathways in the ARC and showing a significant contribution to PCA dimension 2, which discriminates C57BL/6J and WSB/EiJ mice (Figure S2). The statistics analyzing the effects of Strain, Diet and Strain*Diet from qPCR data are also reported for these genes (NS: not significant). Finally, the significant contributions to dimension 2 of the Strain*Diet interaction, the C57BL/6J after 3 days HFD and the WSB/EiJ after 8 weeks of HFD are also provided.

**Table S2**. Summary table of the statistics of the genes involved in inflammation and mitochondrial pathways in the PVN and showing a significant contribution to PCA dimension 2, which discriminates C57BL/6J and WSB/EiJ mice (Figure S2). The statistics analyzing the effects of Strain, Diet and Strain*Diet from qPCR data are also reported for these genes (NS: not significant). Finally, the significant contributions of the Strain*Diet interaction, the C57BL/6J after 3 days HFD and the WSB/EiJ control and after 8 weeks of HFD are also provided.

**Table S3**. List of the genes included in the Heatmap representation and corresponding Taqman primers and probe used for the gene expression analysis for both the arcuate nuclei of the hypothalamus (Figure 7) and the paraventricular nuclei of the hypothalamus (Figure 8).

Figure S1

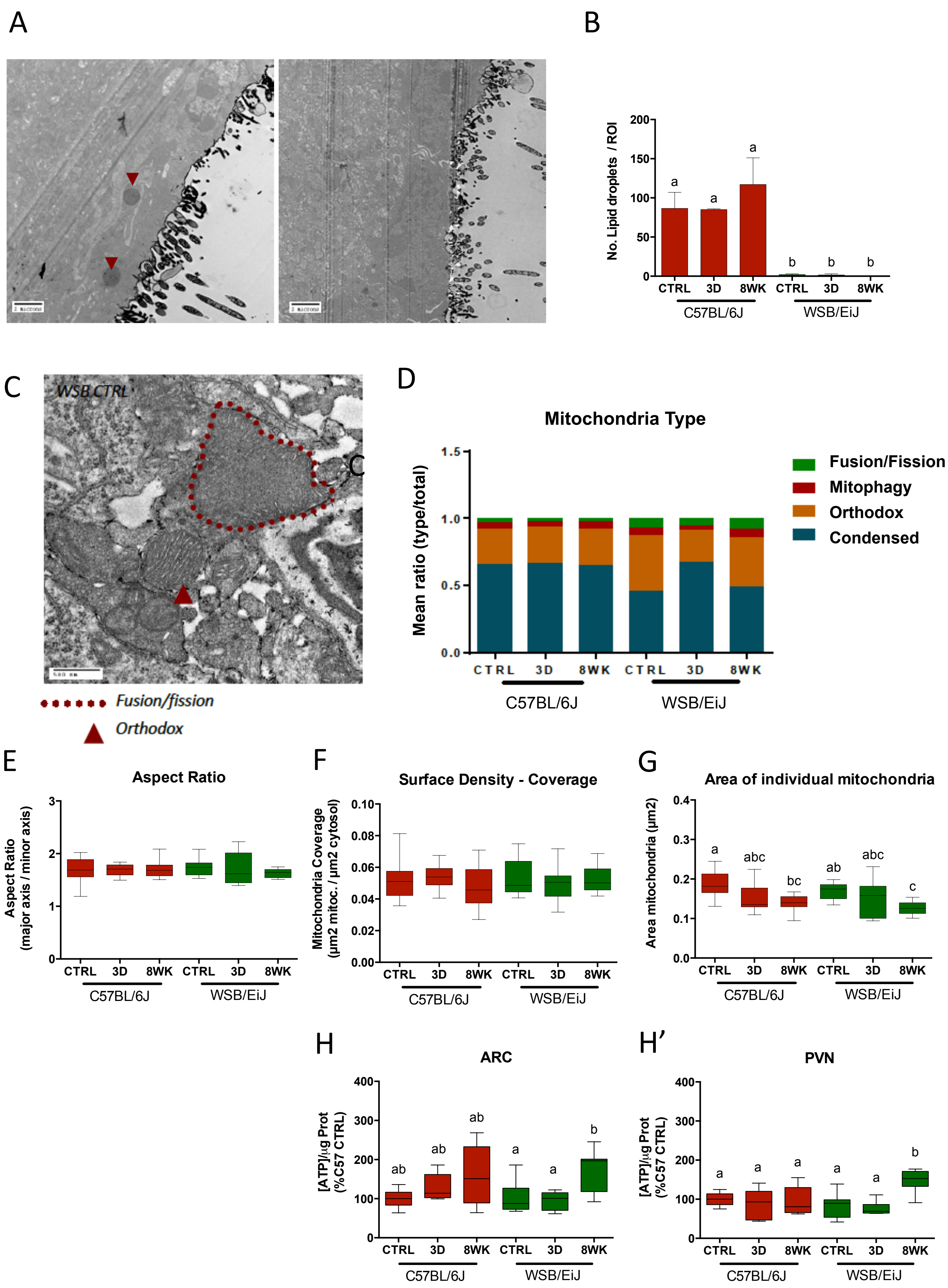
A
B
No. Lipid droplets / ROI
CTRL 3D 8WK CTRL 3D 8WK
C57BL/6J
WSB/EiJ
C
WSB CTRL
Fusion/fission
Orthodox
D
Mitochondria Type
Mean ratio (type/total)
Fusion/Fission
Mitophagy
Orthodox
Condensed
E
Aspect Ratio
Aspect Ratio (major axis / minor axis)
F
Surface Density - Coverage
Mitochondria Coverage (µm2 mitoc. / µm2 cytosol)
G
Area of individual mitochondria
Area mitochondria (µm2)
H
ARC
[ATP]/µg Prot (%C57 CTRL)
H'
PVN

### Figure S2

**ARC**

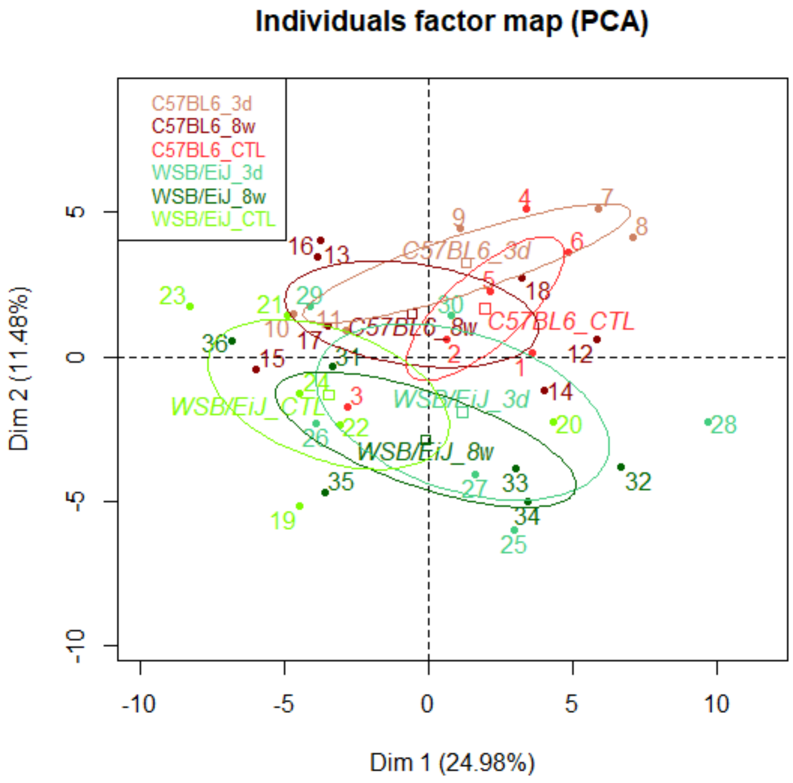


**PVN**

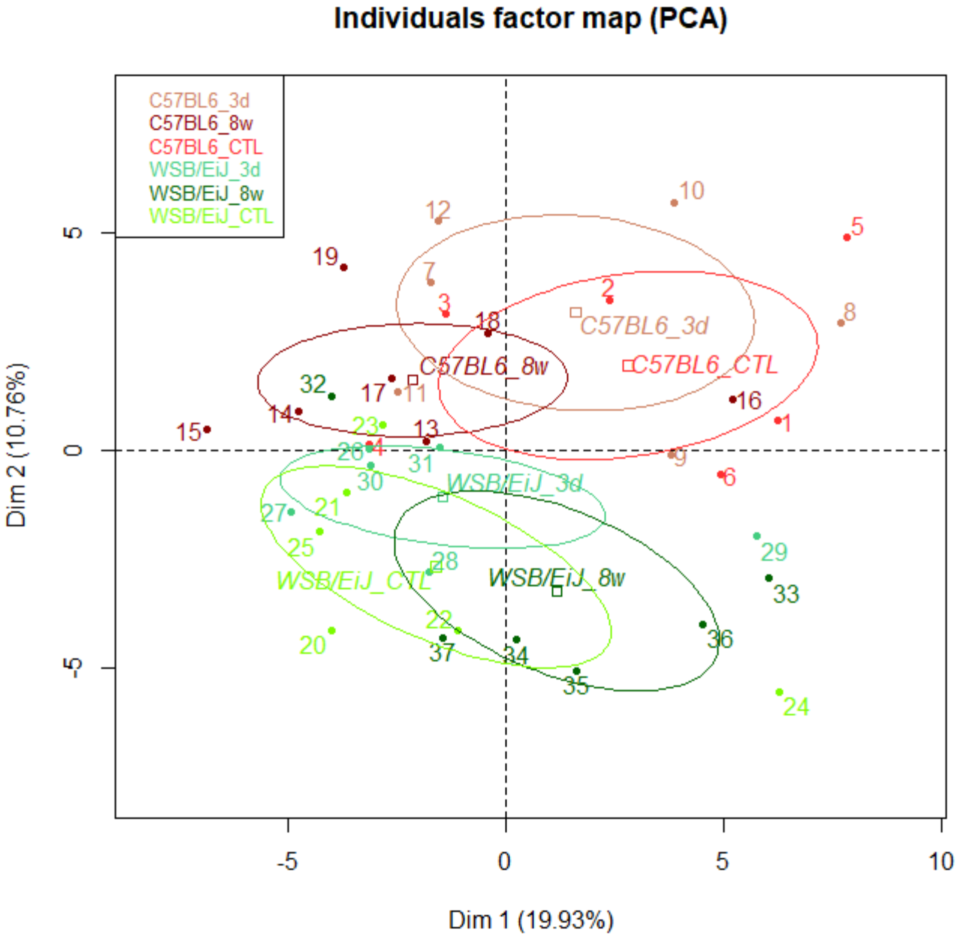

### Figure S3

#### A : ARC : genes more expressed in the WSB/EiJ, related to mitochondria

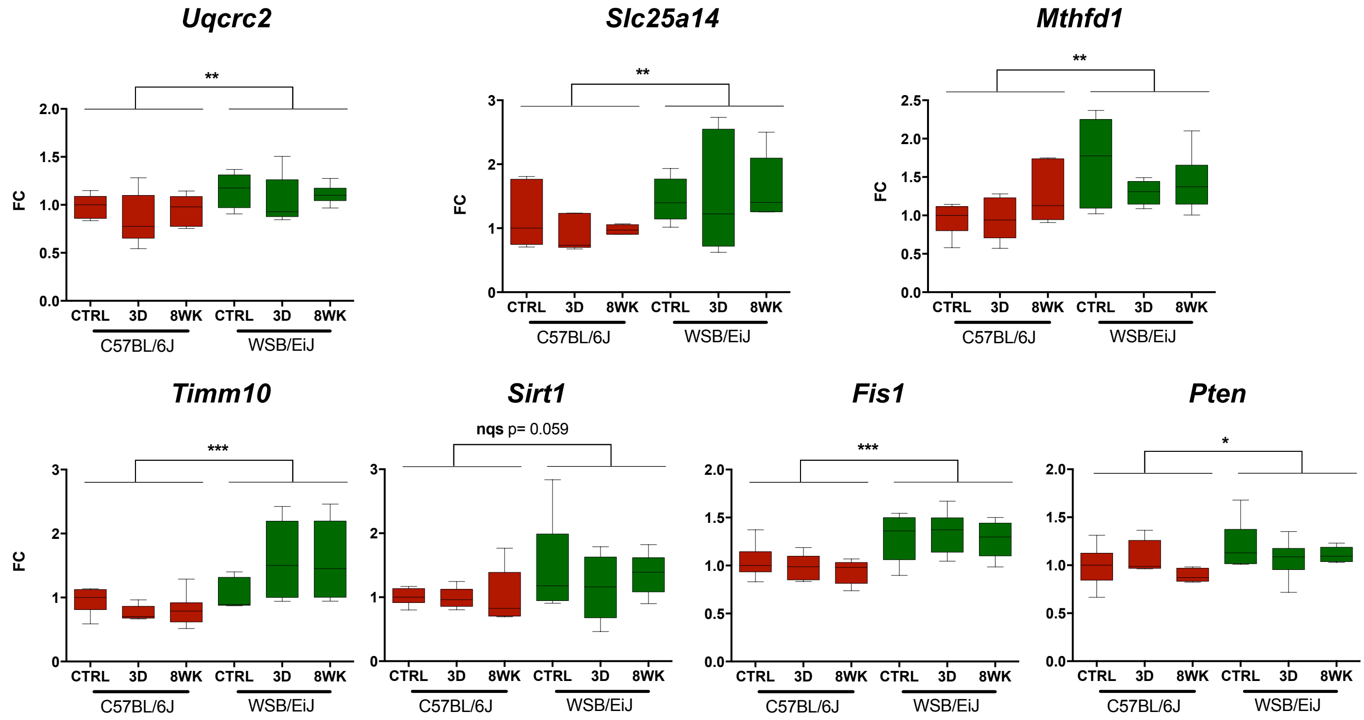


#### B: ARC : genes more expressed in the WSB/EiJ, related to inflammation

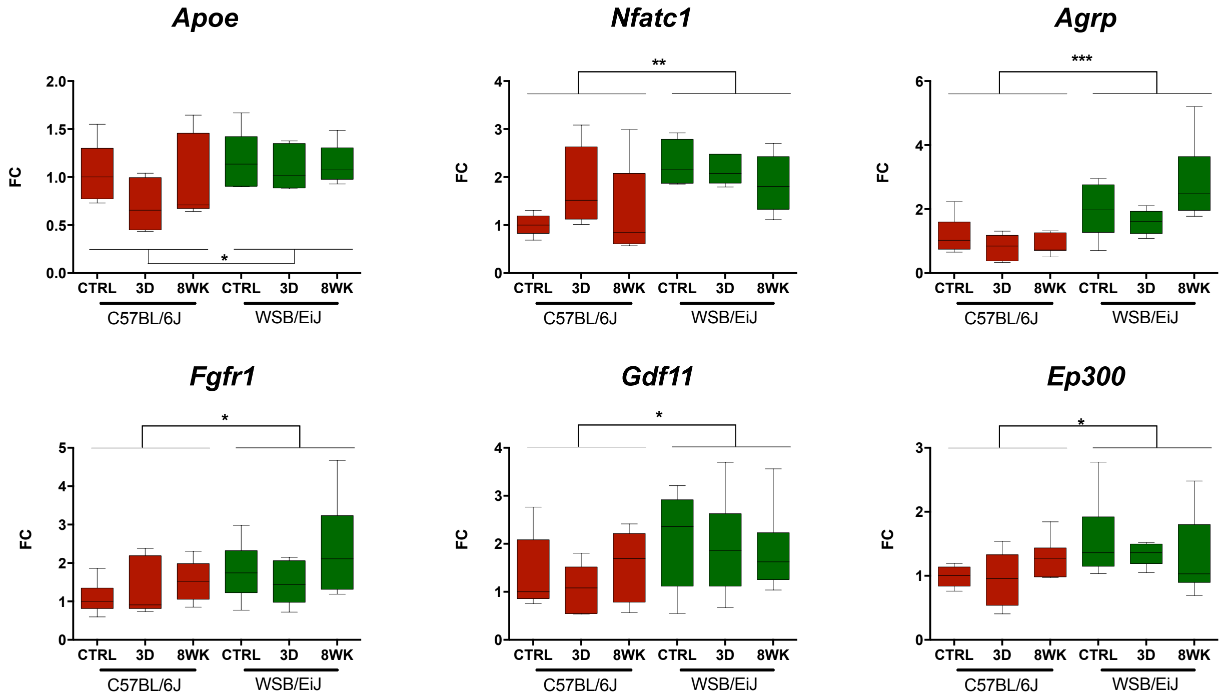

### Figure S4

#### A : PVN: genes more expressed in the WSB/EiJ, related to mitochondria

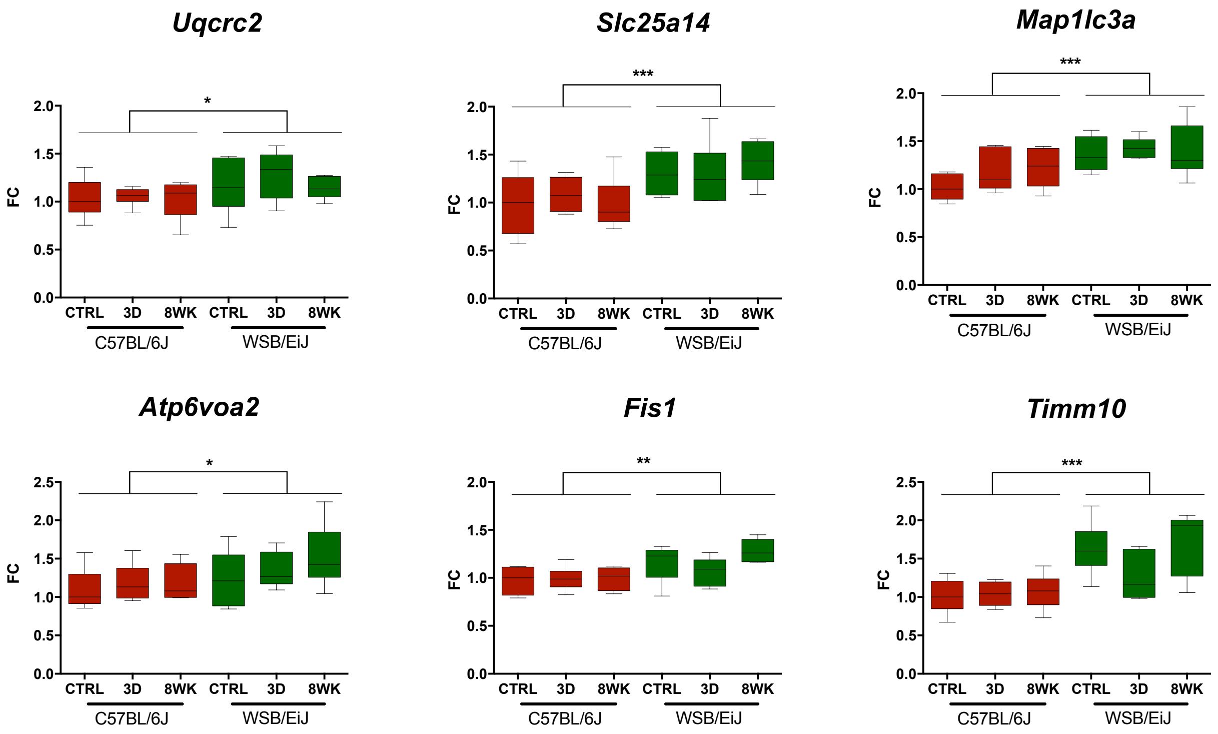


#### B: PVN: genes more expressed in the WSB/EiJ, related to inflammation

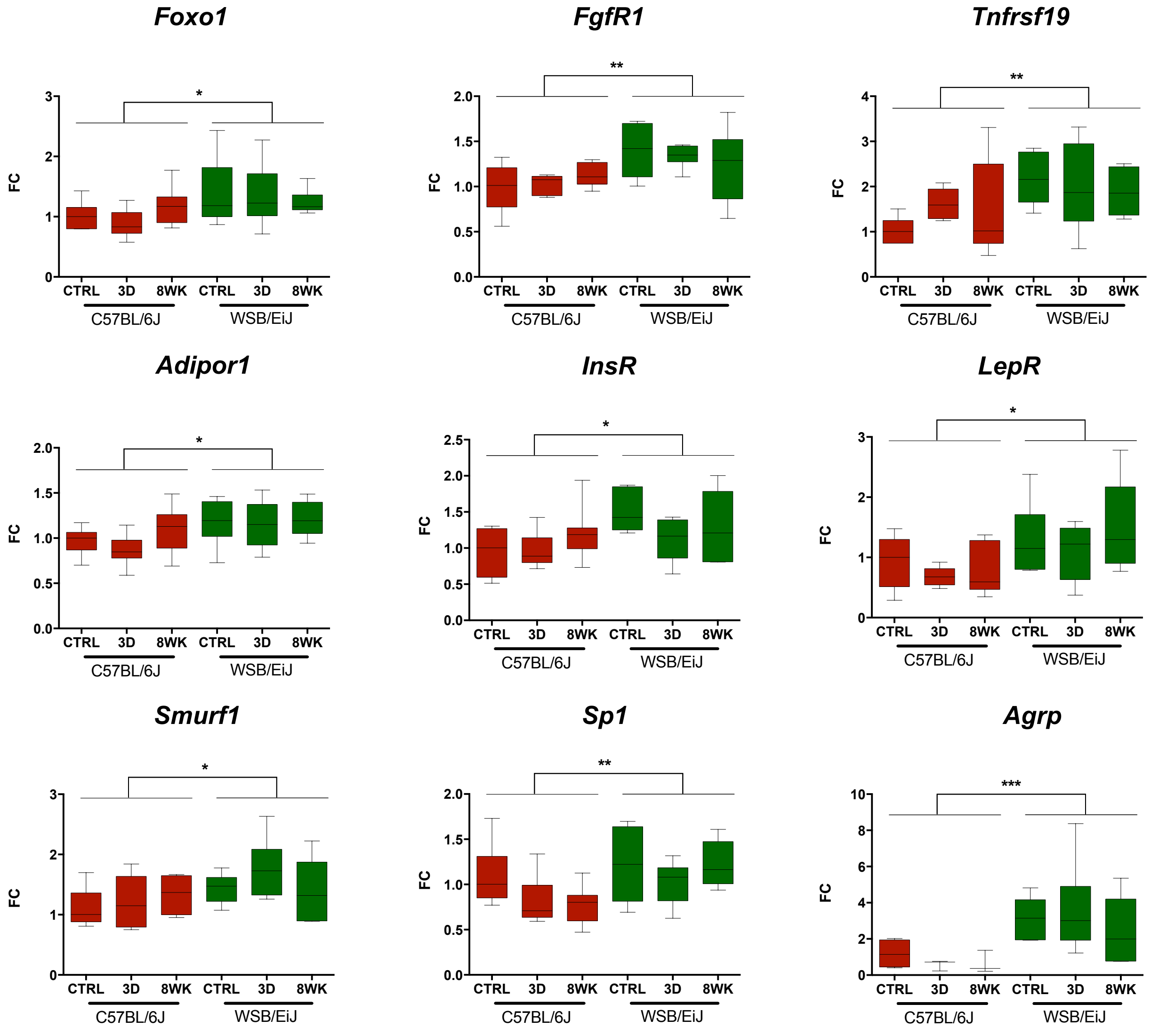

### Figure S5

#### A: ARC : genes less expressed in the WSB/EiJ, related to mitochondria

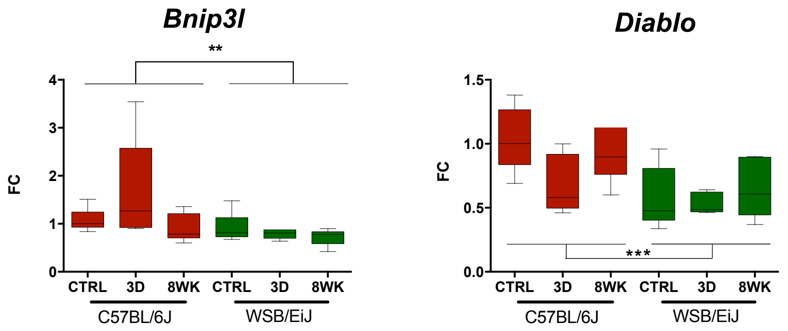


#### B: ARC : genes less expressed in the WSB/EiJ, related to inflammation

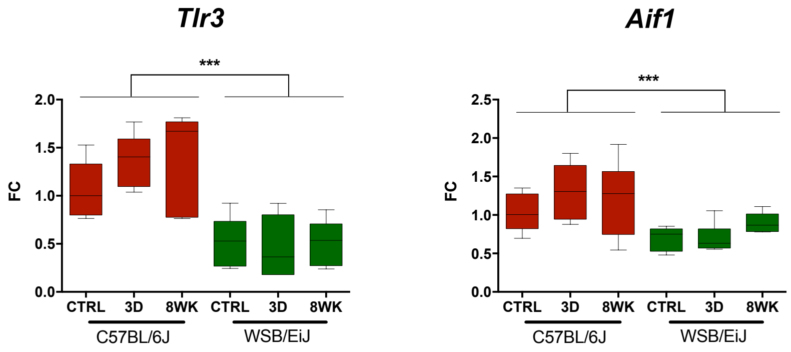


#### C: PVN: genes less expressed in the WSB/EiJ, related to mitochondria

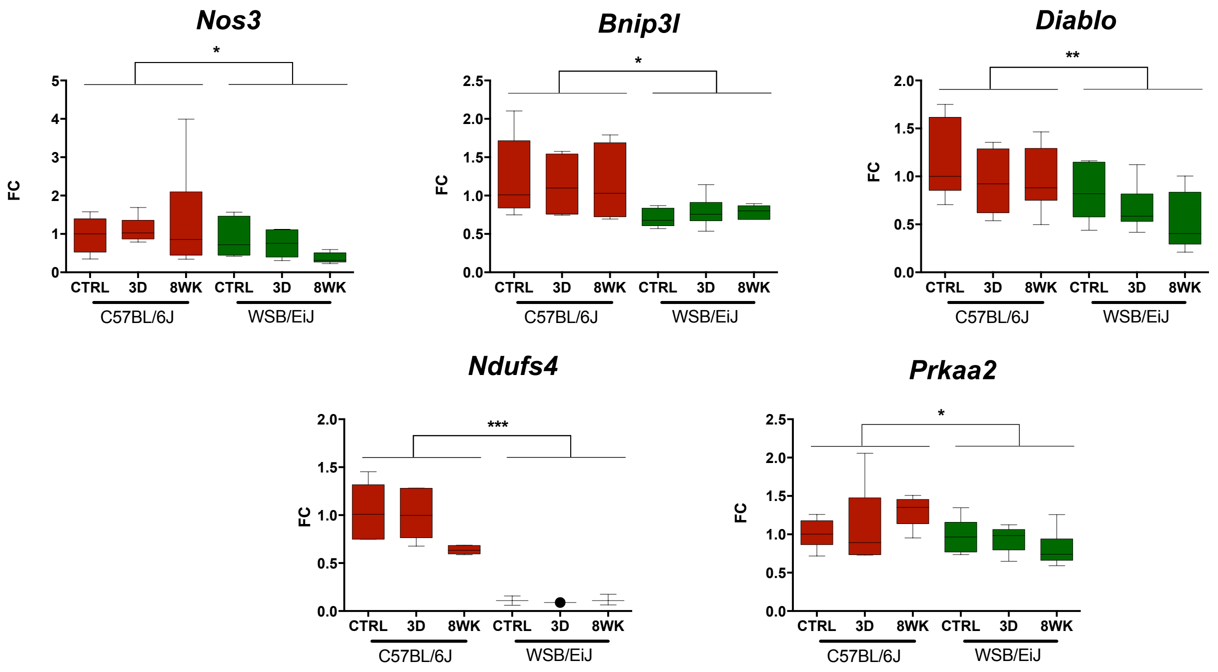


#### D: PVN: genes less expressed in the WSB/EiJ, related to inflammation

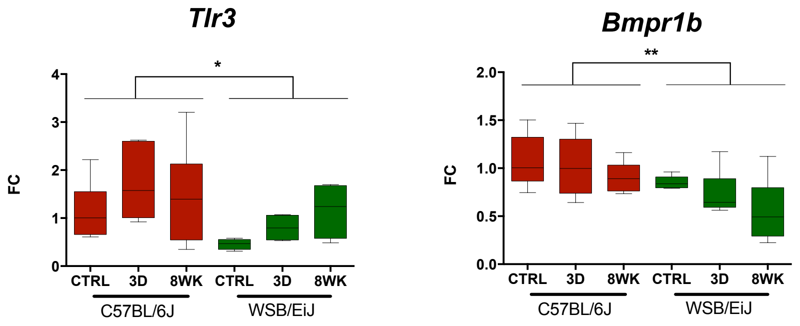

### Figure S6

#### A: ARC : genes differentially regulated by HFD

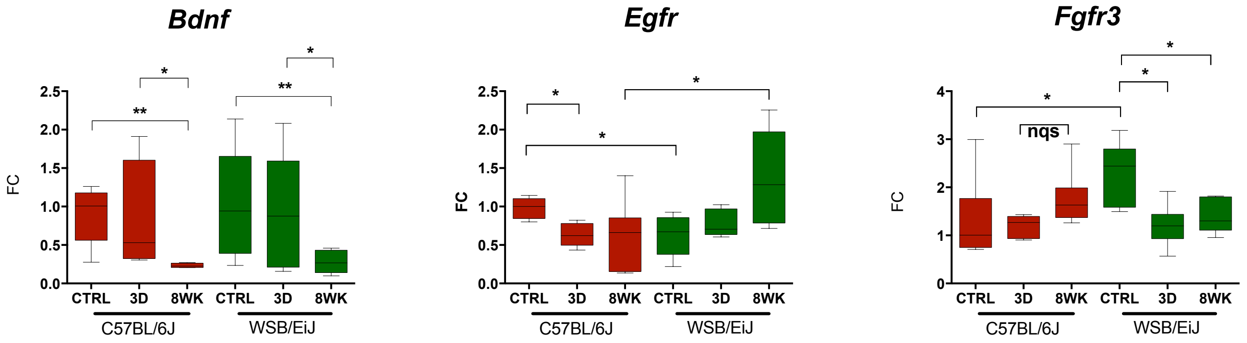


#### B: PVN: genes differentially regulated by HFD, related to Mitochondria

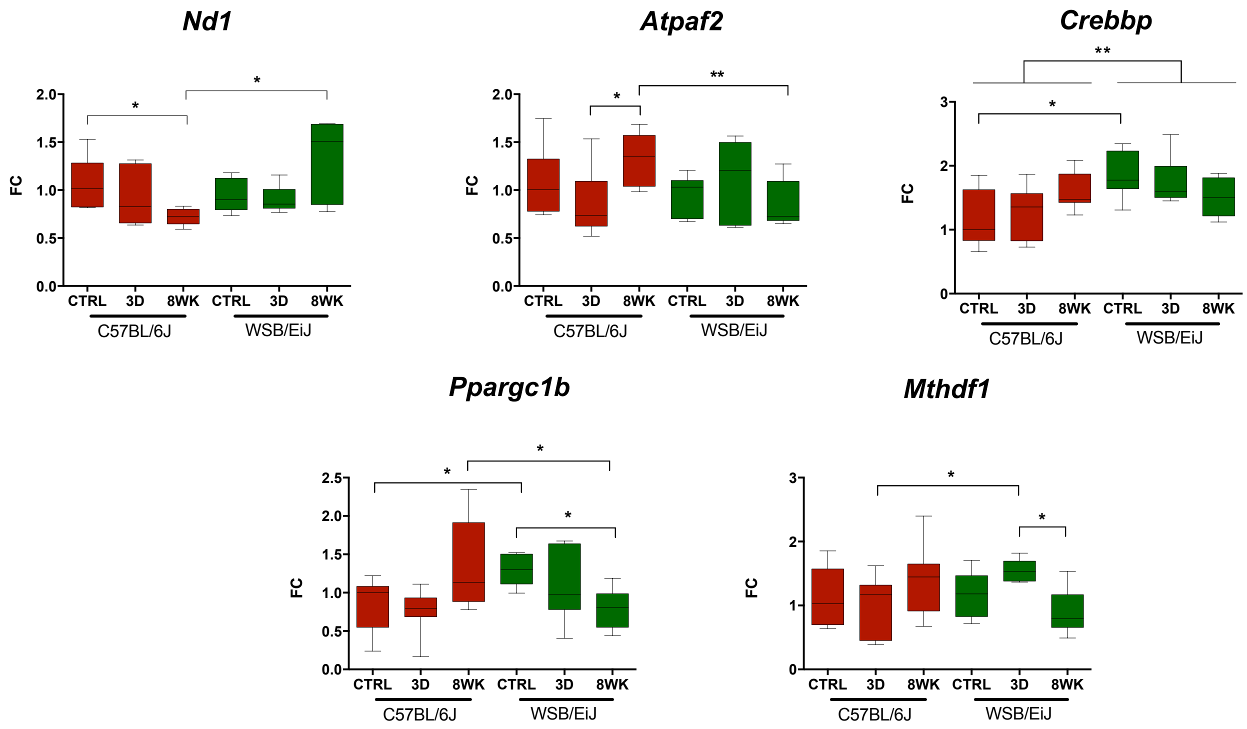


#### C: PVN: genes differentially regulated by HFD, related to Inflammation

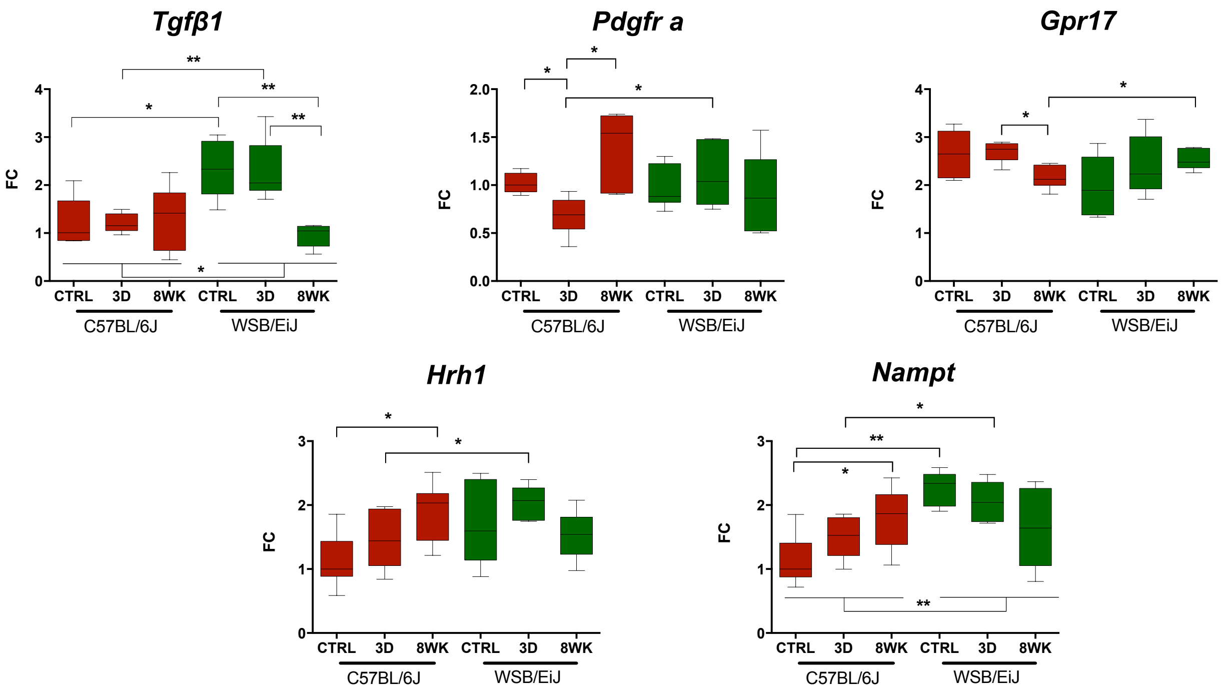

| PCA analysis | | Gene expression : Anova two-way analysis (p-value) | | |
|---|---|---|---|---|
| Inflammatory Genes | p-value | Strain | Diet | Strain*Diet |
| AGRP | 3.834689e-05 | 0.0001 | NS | NS |
| APOE | 8.039549e-05 | 0.01 | NS | NS |
| ADIPOR2 | 3.278060e-03 | NS | NS | NS |
| NFATC1 | 4.390677e-03 | 0.003 | NS | NS |
| LEPR | 6.046190e-03 | NS | NS | NS |
| MTOR | 6.977474e-03 | 0.015 | NS | NS |
| FOXO3 | 9.368151e-03 | NS | NS | NS |
| FLT1 | 1.330187e-02 | NS | NS | NS |
| INSR | 1.332710e-02 | 0.011 | NS | 0.038 |
| CAMKK1 | 1.392742e-02 | NS | NS | NS |
| BMPR1B | 3.138057e-02 | NS | NS | NS |
| NAMPT | 4.301666e-02 | NS | NS | NS |
| TNFRSF19 | 4.683527e-02 | NS | 0.041 | NS |
| AKT1 | 1.394955e-02 | NS | NS | NS |
| TLR3 | 7.134517e-03 | 0.0001 | NS | NS |
| Aif1 | 1.469956e-03 | 0.0003 | NS | NS |

| PCA analysis | | Gene expression : Anova two-way analysis (p-value) | | |
|---|---|---|---|---|
| Mitochondrial Genes | p-value | Strain | Diet | Strain*Diet |
| FIS1 | 6.010403e-09 | 0.0001 | NS | NS |
| SLC25A14 | 4.538424e-07 | 0.001 | NS | NS |
| TIMM10 | 1.076128e-04 | 0.0001 | NS | NS |
| Mthfd1 | 4.928296e-04 | NS | NS | NS |
| COX1 | 1.389519e-03 | NS | NS | NS |
| ND4 | 4.471645e-03 | NS | NS | NS |
| UQCRC2 | 6.150910e-03 | 0.007 | NS | NS |
| SLC25A25 | 1.878896e-02 | NS | NS | NS |
| ND5.315 | 2.226845e-02 | NS | NS | NS |
| NDUFV1 | 2.809522e-02 | NS | NS | NS |
| COX2 | 3.935374e-02 | NS | NS | NS |
| CYTB | 4.761368e-02 | NS | NS | NS |
| NOS3 | 1.526803e-02 | NS | NS | NS |
| BNIP3L | 1.245651e-05 | 0.003 | NS | NS |

| PCA analysis | | Gene expression : Anova two-way analysis (p-value) | | |
|---|---|---|---|---|
| Mitochondrial & Inflammation Genes | p-value | Strain | Diet | Strain*Diet |
| PRKAA1 | 8.953016e-06 | NS | NS | NS |
| SP1 | 1.504849e-03 | NS | NS | NS |
| SIRT1 | 8.080727e-03 | 0.041 | NS | NS |
| PRKAA2 | 4.689706e-02 | NS | NS | NS |

| $Dim.2$ QUALI | | |
|---|---|---|
| | R2 | p.value |
| Strain*Diet | 0.483942 | 0.000896946 |

| $Dim.2$ CATEGORY | | |
|---|---|---|
| | Estimate | p.value |
| C57BL6_3d | 3.190887 | 0.01045966 |
| WSB/EiJ_8w | -2.917066 | 0.01173892 |

**Tableau S1 ARC**

| PCA analysis | | Gene expression : Anova two-way analysis (p-value) | | |
|---|---|---|---|---|
| Inflammatory Genes | p-value | Strain | Diet | Strain*Diet |
| LEPR | 5.736239e-04 | 0.006 | NS | NS |
| FOXO3 | 9.957747e-04 | 0.004 | NS | NS |
| BDNF | 2.201794e-03 | 0.024 | 0.039 | NS |
| INSR | 4.721540e-03 | 0.021 | NS | NS |
| FOXO1 | 6.820992e-03 | 0.013 | NS | NS |
| ADIPOR1 | 8.460876e-03 | 0.004 | NS | NS |
| HIF1A | 1.229405e-02 | NS | NS | NS |
| ADIPOR2 | 3.160265e-02 | NS | NS | NS |
| GDF11 | 4.789413e-02 | NS | NS | NS |
| IRS1 | 4.861798e-02 | NS | NS | NS |
| TLR3 | 5.835337e-03 | 0.006 | NS | NS |
| BMPR1B | 2.054803e-03 | 0.002 | 0.045 | NS |

| PCA analysis | | Gene expression : Anova two-way analysis (p-value) | | |
|---|---|---|---|---|
| **Mitochondrial Genes** | p-value | Strain | Diet | Strain*Diet |
| FIS1 | 2.324651e-05 | 0.002 | NS | NS |
| TIMM10 | 3.138049e-05 | 0.0001 | NS | NS |
| CYTB | 1.443936e-04 | 0.047 | NS | NS |
| ND4 | 1.938533e-04 | NS | NS | NS |
| ND2 | 3.051240e-04 | NS | NS | NS |
| ND5.315 | 5.832932e-04 | NS | NS | NS |
| ND1 | 6.263681e-04 | NS | NS | NS |
| COX2 | 1.427020e-03 | NS | NS | NS |
| COX1 | 4.528668e-03 | NS | NS | NS |
| ND6.325 | 7.060683e-03 | NS | NS | NS |
| SLC25A14 | 1.068456e-02 | 0.002 | NS | NS |
| ATP6V0A2 | 2.916285e-02 | 0.03 | NS | NS |
| MFN1 | 4.983646e-02 | NS | NS | NS |
| NOS3 | 1.237956e-02 | 0.011 | NS | NS |
| DIABLO | 2.205347e-03 | 0.001 | NS | NS |
| NDUFS4 | 1.952163e-04 | 0.0001 | NS | NS |

| PCA analysis | | Gene expression : Anova two-way analysis (p-value) | | |
|---|---|---|---|---|
| **Mitochondrial & Inflammation Genes** | p-value | Strain | Diet | Strain*Diet |
| DNM1L | 2.512209e-03 | NS | NS | NS |
| CREBBP | 3.926364e-03 | 0.003 | NS | 0.032 |
| SP1 | 1.503648e-05 | 0.002 | NS | NS |
| PRKAA1 | 3.859794e-02 | NS | NS | 0.031 |

| $Dim.2$ QUALI | | |
|---|---|---|
| | R2 | p.value |
| Strain*Diet | 0.6414819 | 3.465914e-06 |

| $Dim.2$ CATEGORY | | |
|---|---|---|
| | Estimate | p.value |
| C57BL6_3d | 3.217381 | 0.003453725 |
| WSB/EiJ_CTL | -2.565432 | 0.018532461 |
| WSB/EiJ_8w | -3.239763 | 0.002357155 |

**Tableau S2 : PVN**

**Gene List used for heatmap**

| MGI Gene/Marker ID | Symbol | Name | Entrez Gene ID | Assay ID |
|---|---|---|---|---|
| MGI:87986 | Akt1 | thymoma viral proto-oncogene 1 | 11651 | Mm01331626_m1 |
| MGI:1261423 | Casp8 | caspase 8 | 12370 | Mm00802247_m1 |
| MGI:1097153 | Cx3cl1 | chemokine (C-X3-C motif) ligand 1 | 20312 | Mm00436454_m1 |
| MGI:95522 | Fgfr1 | fibroblast growth factor receptor 1 | 14182 | Mm00438930_m1 |
| MGI:1890077 | Foxo1 | forkhead box O1 | 56458 | Mm00490672_m1 |
| MGI:1861437 | Gsk3b | glycogen synthase kinase 3 beta | 56637 | Mm00444911_m1 |
| MGI:1349419 | Aifm1 | apoptosis-inducing factor, mitochondrion-associated 1 | 26926 | Mm00442540_m1 |
| MGI:88057 | Apoe | apolipoprotein E | 11816 | Mm01307193_g1 |
| MGI:88145 | Bdnf | brain derived neurotrophic factor | 12064 | Mm04230607_s1 |
| MGI:107191 | Bmpr1b | bone morphogenetic protein receptor, type 1B | 12167 | Mm03023971_m1 |
| MGI:1332659 | Bnip3l | BCL2/adenovirus E1B interacting protein 3-like | 12177 | Mm00786306_s1 |
| MGI:1913843 | Diablo | diablo homolog (Drosophila) | 66593 | Mm01194441_m1 |
| MGI:1276116 | Ep300 | E1A binding protein p300 | 328572 | Mm00625535_m1 |
| MGI:1915661 | Map1lc3a | microtubule-associated protein 1 light chain 3 alpha | 66734 | Mm00458724_m1 |
| MGI:1914693 | Map1lc3b | microtubule-associated protein 1 light chain 3 beta | 67443 | Mm00782868_sH |
| MGI:97250 | Myc | myelocytomatosis oncogene | 17869 | Mm00487804_m1 |
| MGI:102469 | Nfatc1 | nuclear factor of activated T cells, cytoplasmic, calcineurin dependent 1 | 18018 | Mm00479445_m1 |
| MGI:97530 | Pdgfra | platelet derived growth factor receptor, alpha polypeptide | 18595 | Mm00440701_m1 |
| MGI:109583 | Pten | phosphatase and tensin homolog | 19211 | Mm00477208_m1 |
| MGI:2135607 | Sirt1 | sirtuin 1 | 93759 | Mm00490758_m1 |
| MGI:107810 | Tfam | transcription factor A, mitochondrial | 21780 | Mm00447485_m1 |
| MGI:98725 | Tgfb1 | transforming growth factor, beta 1 | 21803 | Mm01178820_m1 |
| MGI:2385459 | Socs5 | suppressor of cytokine signaling 5 | 56468 | Mm00465631_s1 |
| MGI:1352474 | Tnfrsf19 | tumor necrosis factor receptor superfamily, member 19 | 29820 | Mm00443506_m1 |
| MGI:1914664 | Mfn1 | mitofusin 1 | 67414 | Mm00612599_m1 |
| MGI:2442230 | Mfn2 | mitofusin 2 | 170731 | Mm00500120_m1 |
| MGI:98372 | Sp1 | trans-acting transcription factor 1 | 20683 | Mm00489039_m1 |
| MGI:104855 | Atp6v0a2 | ATPase, H+ transporting, lysosomal V0 subunit A2 | 21871 | Mm00441838_m1 |
| MGI:2180561 | Atpaf2 | ATP synthase mitochondrial F1 complex assembly factor 2 | 246782 | Mm00520660_m1 |
| MGI:1921256 | Dnm1l | dynamin 1-like | 74006 | Mm01342903_m1 |
| MGI:1913687 | Fis1 | fission 1 (mitochondrial outer membrane) homolog (yeast) | 66437 | Mm00481580_m1 |
| MGI:1338027 | Gdf11 | growth differentiation factor 11 | 14561 | Mm01159973_m1 |
| MGI:102504 | mt-Co1 | mitochondrially encoded cytochrome c oxidase I | 17708 | Mm04225243_g1 |
| MGI:102503 | mt-Co2 | mitochondrially encoded cytochrome c oxidase II | 17709 | Mm03294838_g1 |
| MGI:102501 | mt-Cytb | mitochondrially encoded cytochrome b | 17711 | Mm04225271_g1 |
| MGI:101787 | mt-Nd1 | mitochondrially encoded NADH dehydrogenase 1 | 17716 | Mm04225274_s1 |
| MGI:102500 | mt-Nd2 | mitochondrially encoded NADH dehydrogenase 2 | 17717 | Mm04225288_s1 |
| MGI:102498 | mt-Nd4 | mitochondrially encoded NADH dehydrogenase 4 | 17719 | Mm04225294_s1 |
| MGI:102496 | mt-Nd5 | mitochondrially encoded NADH dehydrogenase 5 | 17721 | Mm04225315_s1 |
| MGI:102495 | mt-Nd6 | mitochondrially encoded NADH dehydrogenase 6 | 17722 | Mm04225325_g1 |
| MGI:1343135 | Ndufs4 | NADH dehydrogenase (ubiquinone) Fe-S protein 4 | 17993 | Mm00656176_m1 |
| MGI:107851 | Ndufv1 | NADH dehydrogenase (ubiquinone) flavoprotein 1 | 17995 | Mm00504941_m1 |
| MGI:1330823 | Slc25a14 | solute carrier family 25 (mitochondrial carrier, brain), member 14 | 20523 | Mm00488302_m1 |
| MGI:1915913 | Slc25a25 | solute carrier family 25 (mitochondrial carrier, phosphate carrier), member 25 | 227731 | Mm00525104_m1 |
| MGI:1921261 | Slc25a27 | solute carrier family 25, member 27 | 74011 | Mm00511820_m1 |
| MGI:98443 | Surf1 | surfeit gene 1 | 20930 | Mm00489041_g1 |
| MGI:1353429 | Timm10 | translocase of inner mitochondrial membrane 10 | 30059 | Mm00443538_m1 |
| MGI:1914253 | Uqcrc2 | ubiquinol cytochrome c reductase core protein 2 | 67003 | Mm00445961_m1 |
| MGI:1919924 | Adipor1 | adiponectin receptor 1 | 72674 | Mm01291334_mH |
| MGI:892013 | Agrp | agouti related neuropeptide | 11604 | Mm00475829_g1 |
| MGI:1890081 | Foxo3 | forkhead box O3 | 56484 | Mm01185722_m1 |
| MGI:1338071 | Ikbkb | inhibitor of kappaB kinase beta | 16150 | Mm01222247_m1 |
| MGI:96575 | Insr | insulin receptor | 16337 | Mm01211875_m1 |
| MGI:99454 | Irs1 | insulin receptor substrate 1 | 16367 | Mm01278327_m1 |
| MGI:104993 | Lepr | leptin receptor | 16847 | Mm00440181_m1 |
| MGI:1928394 | Mtor | mechanistic target of rapamycin (serine/threonine kinase) | 56717 | Mm00444968_m1 |
| MGI:2145955 | Prkaa1 | protein kinase, AMP-activated, alpha 1 catalytic subunit | 105787 | Mm01296700_m1 |
| MGI:1336173 | Prkaa2 | protein kinase, AMP-activated, alpha 2 catalytic subunit | 108079 | Mm01264789_m1 |
| MGI:109354 | Ucp2 | uncoupling protein 2 (mitochondrial, proton carrier) | 22228 | Mm00627599_m1 |
| MGI:2140940 | Acacb | acetyl-Coenzyme A carboxylase beta | 100705 | Mm01204667_m1 |
| MGI:93830 | Adipor2 | adiponectin receptor 2 | 68465 | Mm01184032_m1 |
| MGI:1891766 | Camkk1 | calcium/calmodulin-dependent protein kinase kinase 1, alpha | 55984 | Mm00517053_m1 |
| MGI:1098296 | Cpt1a | carnitine palmitoyltransferase 1a, liver | 12894 | Mm01231183_m1 |
| MGI:1270849 | Rps6kb1 | ribosomal protein S6 kinase, polypeptide 1 | 72508 | Mm01310033_m1 |
| MGI:1914930 | Sdhb | succinate dehydrogenase complex, subunit B, iron sulfur (Ip) | 67680 | Mm00458272_m1 |
| MGI:1098280 | Crebbp | CREB binding protein | 12914 | Mm01342452_m1 |
| MGI:2444934 | Ppargc1b | peroxisome proliferative activated receptor, gamma, coactivator 1 beta | 170826 | Mm00504720_m1 |
| MGI:107619 | Hrh1 | histamine receptor H1 | 15465 | Mm00434002_s1 |
| MGI:104741 | Nfkbia | nuclear factor of kappa light polypeptide gene enhancer in B cells inhibitor, alpha | 18035 | Mm00477800_g1 |
| MGI:1343098 | Aif1 | allograft inflammatory factor 1 | 11629 | Mm00479862_g1 |
| MGI:1339753 | Csf1 | colony stimulating factor 1 (macrophage) | 12977 | Mm00432686_m1 |
| MGI:95294 | Egfr | epidermal growth factor receptor | 13649 | Mm00433023_m1 |
| MGI:95558 | Flt1 | FMS-like tyrosine kinase 1 | 14254 | Mm00438980_m1 |
| MGI:106918 | Hif1a | hypoxia inducible factor 1, alpha subunit | 15251 | Mm00468869_m1 |
| MGI:1929865 | Nampt | nicotinamide phosphoribosyltransferase | 59027 | Mm00451938_m1 |
| MGI:97362 | Nos3 | nitric oxide synthase 3, endothelial cell | 18127 | Mm00435217_m1 |
| MGI:103038 | Stat3 | signal transducer and activator of transcription 3 | 20848 | Mm01219775_m1 |
| MGI:95524 | Fgfr3 | fibroblast growth factor receptor 3 | 14184 | Mm00433294_m1 |
| MGI:3584514 | Gpr17 | G protein-coupled receptor 17 | 574402 | Mm02619401_s1 |
| MGI:1342005 | Mthfd1 | methylenetetrahydrofolate dehydrogenase (NADP+ dependent), methenyltetrahydrofolate cyclohydrolase, formyltetrahydrofolate synthase | 108156 | Mm00507092_m1 |
| MGI:1338850 | Mthfd2 | methylenetetrahydrofolate dehydrogenase (NAD+ dependent), methenyltetrahydrofolate cyclohydrolase | 17768 | Mm00485276_m1 |
| MGI:104752 | Nfkbib | nuclear factor of kappa light polypeptide gene enhancer in B cells inhibitor, beta | 18036 | Mm00456849_m1 |
| MGI:1921620 | Rptor | regulatory associated protein of MTOR, complex 1 | 74370 | Mm00712676_m1 |
| MGI:1923038 | Smurf1 | SMAD specific E3 ubiquitin protein ligase 1 | 75788 | Mm00547102_m1 |
| MGI:2156367 | Tlr3 | toll-like receptor 3 | 142980 | Mm01207404_m1 |
| MGI:96392 | Icam1 | intercellular adhesion molecule 1 | 15894 | Mm00516023_m1 |
| MGI:2661364 | Neu4 | sialidase 4 | 241159 | Mm00620597_m1 |
| MGI:87904 | Actb | actin, beta | 11461 | Mm00607939_s1 |
| MGI:88127 | B2m | beta-2 microglobulin | 12010 | Mm00437762_m1 |
| MGI:88261 | Canx | calnexin | 12330 | Mm00500330_m1 |
| MGI:88276 | Ctnnb1 | catenin (cadherin associated protein), beta 1 | 12387 | Mm00483033_m1 |
| MGI:95640 | Gapdh | glyceraldehyde-3-phosphate dehydrogenase | 14433 | Mm99999915_g1 |
| MGI:96217 | Hprt | hypoxanthine guanine phosphoribosyl transferase | 15452 | Mm00446968_m1 |
| MGI:97555 | Pgk1 | phosphoglycerate kinase 1 | 18655 | Mm00435617_m1 |
| MGI:98889 | Ubc | ubiquitin C | 22190 | Mm02525934_g1 |
| MGI:109484 | Ywhaz | tyrosine 3-monooxygenase/tryptophan 5-monooxygenase activation protein, zeta polypeptide | 22631 | Mm03950126_s1 |
